# Epithelial-mesenchymal plasticity determines estrogen receptor positive (ER+) breast cancer dormancy and reacquisition of an epithelial state drives awakening

**DOI:** 10.1101/2021.07.22.453458

**Authors:** Patrick Aouad, Yueyun Zhang, Céline Stibolt, Sendurai A. Mani, George Sflomos, Cathrin Brisken

## Abstract

Estrogen receptor α-positive (ER+) breast cancers (BCs) represent more than 70% of all breast cancers and pose a particular clinical challenge because they recur up to decades after initial diagnosis and treatment. The mechanisms governing tumor cell dormancy and latent disease remain elusive due to a lack of adequate models. Here, we compare tumor progression of ER+ and triple-negative (TN) BC subtypes with a clinically relevant mouse intraductal xenografting approach (MIND). Both ER+ and TN BC cells disseminate already during the *in situ*stage. However, TN disseminated tumor cells (DTCs) proliferate at the same rate as cells at the primary site and give rise to macro-metastases. ER+ DTCs have low proliferative indices, form only micro-metastases and lose epithelial characteristics. Expression of *CDH1* is decreased whereas the mesenchymal marker *VIM* and the transcription factors, *ZEB1*/*ZEB2,* which control epithelial-mesenchymal plasticity (EMP) are increased. EMP is not detected earlier during ER+ BC development and not required for invasion or metastasis. *In vivo*, forced transition to the epithelial state through ectopic E-cadherin expression overcomes dormancy with increased growth of lung metastases. We conclude that EMP is essential for the generation of a dormant cell state and the development of latent disease. Targeting exit from EMP is of therapeutic potential.

## Introduction

Breast cancer (BC) is the most commonly diagnosed malignancy worldwide (Sung et al., 2021). Clinical management of BC relies on grade, stage and tumor subtype which can be defined by histology and immunohistochemistry (IHC) and/or molecular profiling by global gene expression or PAM50. Nineteen specific histological subtypes are distinguished, and the remainder are classified as of no special type (NST) (Tan et al., 2020). More than 70% of all BC cases are estrogen receptor positive (ER+) as determined by IHC and are of luminal A or B molecular subtype. ER+ BCs tend to be of lower grade and to have lower proliferative indices than HER2+ and ER-/PR-/HER2-, triple negative (TN) BCs as evaluated by Ki67 IHC. Accordingly, patients with ER+ disease have better 5-year survival rates than patients with HER2+ or TN BCs. However, patients with ER- BC who respond to chemotherapy and do not relapse within the first 5 years after treatment are generally considered disease-free. This is not the case for ER+ BC patients; they benefit from endocrine therapy but remain life long at risk with stable recurrence rates (Pan et al., 2017).

Delayed recurrence is attributed to disseminated tumor cells (DTCs) which leave the primary tumor early during tumorigenesis and remain dormant at distant sites (Klein, 2011). The existence of such dormant DTCs is supported by the clinical observation that cancer transmission occurred from organ transplantation from patients with undiagnosed BC and resulted in ER+ adenocarcinoma lesions in the recipients (Buell et al., 2004; Matser et al., 2018). This phenomenon has also been observed in other tumor types such as melanoma, and prostate cancer (Crowley and Seigler, 1990; Strauss and Thomas, 2010; Tsao et al., 1997; Weckermann et al., 2001). It is critical to understand the mechanisms governing dormancy for preventative intervention, however, the field has been severely understudied due to the lack of suitable in vivo models (reviewed by Klein, 2020; Richman and Dowsett, 2019; Risson et al., 2020; Zhang et al., 2013). Most studies of the mechanisms that underly metastasis *in vivo* are conducted using genetically engineered mouse models (GEMMs). With regards to mammary tumorigenesis, these do not recapitulate the subtype diversity of the human disease as the resulting mammary carcinomas are mostly ER- (Green et al., 2000; Guy et al., 1992; Sinn et al., 1987) whilst dormancy is a characteristic of ER+ BC. Additionally, most xenograft studies have relied on subcutaneous grafting of TN cell line models or injection of BC cell lines, including ER+ cells into the bloodstream or at distant sites, which poorly reflect the clinical situation.

We recently reported that grafting of human ER+ BC cell lines and patient-derived xenografts (PDXs), to the milk ducts of immune-compromised mice (MIND) enables their *in vivo* growth in a physiological endocrine milieu and substantially improves take rates over the traditional subcutaneous engraftment site (Fiche et al., 2019; Sflomos et al., 2016). The same applies to molecular apocrine breast cancers (Richard et al., 2016). Such ER+ intraductal xenograft models recapitulate human disease progression from *in situ*stage to spontaneous dissemination to clinically relevant distant organs (Fiche et al., 2019; Sflomos et al., 2016) and reflect specific histopathological subtypes such as the invasive lobular carcinoma (ILC) (Kozma et al., 2021; Sflomos et al., 2021). Bioluminescence emanating from ER+ DTCs can be detected in different organs when tumor cells still appear to be confined within the milk ducts suggesting early dissemination of DTCs from the primary tumor (Fiche et al., 2019; Sflomos et al., 2016). While metastatic load increases with prolonged *in vivo* tumor growth no macro-metastases are detected up to the experimental endpoint, i.e. 7 months after engraftment of MCF-7 cells (Sflomos et al., 2016) and >1 year after engraftment of ER+ PDXs (Fiche et al., 2019).

Here, we compare ER+ and TNBC progression in MIND models and show that dormancy is specific for ER+ DTCs, and is driven by Epithelial-Mesenchymal Plasticity (EMP). Forced exit from EMP through E-cadherin (E-cad) overexpression can overcome dormancy providing a targetable mechanism against latent disease.

## Significance

Late recurrence of ER+ BC is a major clinical problem. Currently, most experimental studies rely on ER- models and on primary tumor growth as functional readout, we present xenograft models of the clinically relevant ER+ latent disease. Intraductal engraftment of BC cells reveal that in TN BC, DTCs proliferate whereas ER+ DTCs enter dormancy determined by EMP. Conversion of dormant DTCs to an epithelial state leads to recurrence providing a targetable mechanism for therapeutic intervention.

## Results

### Tumor growth and metastatic progression in ER+ *versus* TN BC intraductal xenografts

We observed that ER+ BC cells that are xenografted to milk ducts of immunocompromised mice recapitulate disease progression from the *in situ*stage to dissemination to distant organs (Sflomos et al., 2016). However, during the experimental life span of the host animals, DTCs detected in clinically relevant distant organs failed to progress to macro-metastases. To test whether the ER+ BC MIND models reflect tumor dormancy specific to ER+ BCs, we compared the metastatic behavior of ER+ and ER- BC cells by this aproach. We modelled ER- BC using the TNBC cell lines, BT20 and HCC1806 (Supplementary Table 1), as well as a PDX derived from an untreated primary TNBC, called T70 (Supplementary Table 2). To model ER+ BC, we used the established ER+ cell lines, MCF-7 and T47D (Supplementary Table 1) and 2 ER+ PR+ HER2-PDXs. One of them, T99, was derived from an untreated primary tumor and the other, METS15, from ascites of a patient with advanced disease (Supplementary Table 2). All tumor cells were infected with *GFP:luc* or *RFP:luc* expressing lentiviruses and injected to the milk ducts of NSG females (Fig. 1a) (Sflomos et al., 2016). Take rates, defined as the percentage of glands showing *in vivo* tumor cell growth relative to the total number of glands injected, were 100% for TN and ≥ 90% for ER+ BC xenografts (Extended Data Fig. 1a). Within 5 weeks, *in vivo* bioluminescence of BT20, HCC1806, and T70 xenografts increased 702-, 592-, and 1915-fold, respectively (Fig. 1b, and Extended Data Fig. 1b). Mammary tumors were palpable, the host mice showed signs of morbidity and were euthanized. Over the same time, bioluminescence from MCF-7, T47D, T99, and METS15 xenografts increased only 42-, 39-, 54-, and 5-fold, respectively (Fig. 1b, and Extended Data Fig. 1b). Five months later, bioluminescence increased 1000- to 2000-fold reative to the start and tumors became palpable while the host mice remained healthyy (Fig. 1b, Extended Data Fig. 1c).

**Fig. 1.**
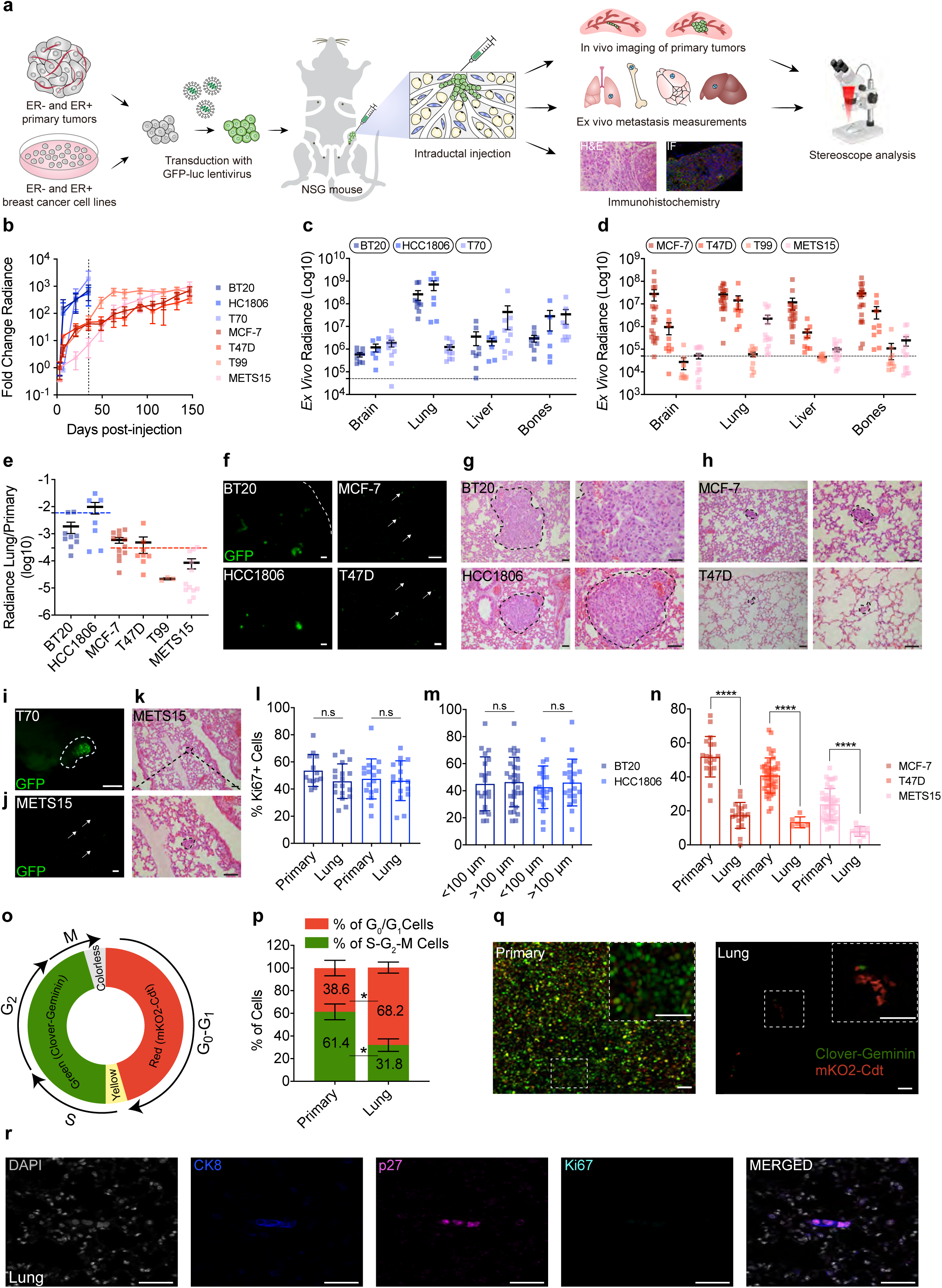
Intraductal xenografts of ER+ and ER- BC cells show distinct growth and metastatic behavior. **a**. Scheme illustrating the intraductal xenograftig approach used in this study. **b.** Graph showing the fold-change of bioluminescence over time for all intraductal xenografts. Data represent mean ± SEM of 13-19 mammary glands from 5-9 mice bearing BT20, HCC1806, T70, MCF-7, T47D, T99, or METS15 xenografts, respectively. The dashed line represents the experimental end-point for TNBC xenografts. **c.** Plot showing ex vivo bioluminescence of resected organs from 9, 7, and 6 mice bearing BT20, HCC1806, and T70 xenografts, respectively. Data represent mean ± SEM. **d.** Plot showing ex vivo bioluminescence on resected organs from 20, 9, 10, and 14 mice bearing MCF-7, T47D, T99, and METS15 xenografts, respectively. Data represent mean ± SEM. **e.** Plot showing the ratio of bioluminescence in lungs over primary tumor. Data represent mean ± SEM. **f.** Representative fluorescence stereo micrographs of lungs from at least 3 mice bearing BT20, HCC1806, MCF-7, and T47D intraductal xenografts. Arrows point to DTCs in ER+ intraductal xenografts-bearing mice. Scale bar, 1 mm. **g.** Representative H&E micrographs of lung sections from 3 mice bearing BT20 and HCC1806, and **h.** MCF-7, and T47D, intraductal xenografts. Scale bar, 50 µm. **i**. Representative fluorescence stereo micrographs of the liver and **j.** lungs from at least 3 mice bearing T70 and METS15 intraductal xenografts, respectively. Scale bars, 1 mm. **k.** Representative H&E micrographs of lung sections from 3 mice bearing METS15 intraductal xenografts. Scale bars, 50 µm. **l.** Percentage of Ki67+ cells in matched primary and lung sections from BT20 and HCC1806 intraductal xenografts 5 weeks after intraductal injection. Data represent mean ± SD from 3 different hosts. Student’s unpaired t-test, two-tailed. **m.** Bar graph showing Ki67 index of small (<100 μm) and large (>100 μm) BT20 and HCC1806 lung lesions. Data represent mean ± SD from 3 different hosts. Student’s unpaired t-test, two-tailed. Each data point in the primary tumor and the lung represents ≥ 1,000 and 100 cells analyzed, respectively. **n.** Percentage of Ki67+ cells in matched primary and lung sections from MCF-7, T47D, and METS15 intraductal xenografts 5 months after intraductal injection. Data represent mean ± SD from 3 different hosts. Student’s paired t-test, two-tailed. **o.** Scheme showing the FUCCI cell cycle reporter. **p.** Bar plot showing the percentage of cycling and non-cycling cells in primary tumors and lung micro-metastases in MCF-7 intraductal xenografts-bearing mice. Data represent mean ± SD from 3 host mice. Paired t-test. **q.** Representative fluorescence micrographs of red, green, and double-positive MCF-7:FUCCI reporter cells in matched primary tumor (left) and lung (right). Scale bars, 50 µm. **r.** Representative immunofluorescence micrographs for CK8 (blue), p27 (magenta) and Ki67 (cyan), counterstained with DAPI (blue) on lung section from MCF-7-bearing mice. Arrows point to p27+/Ki67-lung DTCs. Scale bar, 50 µm. *, ***, ****, and n.s represent *P*<0.05, 0.001, 0.0001, and not significant, respectively.

Ex vivo imaging of resected organs from ER+ and TN models revealed bioluminescence in the brain, lungs, liver and bones (Fig. 1c, d). The signal attributable to BT20 and HCC1806 cells was particularly high in the lungs at 10^7^-10^9^ p/sec/cm^2^/sr (Fig. 1c). T70 cells, derived from a patient with pregnancy-associated BC were most readily detected in the liver with 10^6^-10^8^ p/sec/cm^2^/sr and to lesser extent in the lungs (Fig. 1c) in line with clinical reports (Goddard et al., 2017). Bioluminescence in the lungs and bones resected from MCF-7 and T47D-bearing mice, ranged between 10^6^ and 10^7^ p/sec/cm^2^/sr and the signal attributable to disseminated METS15 cells was 10^5^-10^6^ p/sec/cm^2^/sr (Fig. 1d). PDX-T99 xenografts resulted in bioluminescence just above background levels in the lungs and bones in 3 out of 10 mice (Fig. 1d) consistent with previous observations (Fiche et al., 2019). At the respective experimental endpoints, primary tumor burden was comparable but bioluminescence of the lungs relative to that of the mammary glands, was on average 100-fold higher in the TN than in the ER+ BC xenograft models (Fig. 1e). Thus, lung metastatic disease in the TN BC xenografts advances faster than in the ER+ xenografts reflecting the BC subtype-specific biology of metastasis.

We then examined early tumor cell seeding by fluorescence stereo microscopy of mammary glands xenografted with BT20:*Luc2-GFP* and HCC1806:*Luc2-GFP* cells. One week after intraductal injection, the GFP signal was confined to the ducts (Extended Data Fig. 1d,e) and histological examination confirmed that tumor cells were within the ducts as characteristic of the *in situ*stage (Extended Data Fig. 1f,g). No disruption of the epithelial-stromal border was detected but the basal membrane and the extracellular matrix (ECM) around mouse ducts filled with tumor cells were thicker than around those devoid of human cells (Extended Data Fig. 1f,g). At this stage, *ex vivo* bioluminescence of the lungs was above background levels, in 5 of 10 BT20- and 2 of 4 HCC1806-intraductal xenograft-bearing mice (Extended Data Fig. 1h). Two weeks after engraftment, multiple invasive foci were detected in the engrafted mammary glands (Extended Data Fig. 1d-g). In all BT20 and HCC1806-intraductal xenograft-bearing mice, the lungs showed increased bioluminescence (Extended Data Fig. 1h) and GFP+ lesions were detected by fluorescence stereo microscopy (Extended Data Fig. 1i). Thus, both TN and ER+ BC intraductal xenografts disseminate during the *in situ* stage as observed in clinical studies (Hüsemann et al., 2008; Sänger et al., 2011). However, DTCs from ER- BCs progress to macro-metastases whereas ER+ DTCs fail to grow to detectable lesions.

### ER+ metastatic lesions are dormant

The slow increase in bioluminescence at the distant sites in the ER+ xenograft models suggested that the distant lesions are dormant. To test this, we analyzed DTCs in the lungs by fluorescence stereo microscopy and subsequent histological analysis. Both BT20 and HCC1806 cells gave rise to lesions that were readily detected by fluorescence stereo microscopy and histological examination and measured 0.5-3.0 mm in diameter (Fig. 1f, g). In contrast, lung lesions arising from MCF-7 and T47D intraductal xenografts were barely detected by fluorescence stereo microscopy and measured 10-100 µm in diameter in histological sections (Fig. 1f,h). In the case of the TN PDXs, liver lesions were 0.5-1.0 mm in diameter (Fig. 1i). The ER+ METS15 cells were detected by both approaches (Fig. 1j,k), but T99 cells were neither detected by fluorescence stereo microscopy nor by subsequent screening of >40 histological sections from lung lobes of 3 host mice in agreement with the low bioluminescence (10^5^ p/sec/cm^2^/sr) (Fig. 1n).

Next, we quantified the proliferative index of primary and metastatic lesions from TN and ER+ xenografts using Ki67 IHC (Supplementary Table 3). The Ki67 indices for BT20 and HCC1806 primary tumors were 54.7% and 47.5%, respectively and they were comparable to the values for lung metastases (Fig. 1l, Extended Data Fig. 1j), independent of the size of the lesions (<100 μm or large >100 μm) (Fig. 1m). In contrast, the Ki67+ indices in ER+ intraductal primary tumors were on average 3-times higher than in the distant lesions, with values of 51.8% *vs* 17.4% in MCF-7, 40.8% *vs* 13.3% in T47D, and 23.8% *vs* 7.9% in METS15 xenografts-bearing mice, respectively (Fig. 1n and Extended Data Fig. 1k).

To further characterize cell cycle dynamics, we used a fluorescent ubiquitination-based cell cycle indicator (FUCCI) (Bajar et al., 2016), which distinguishes non-cycling cells (G_0_/G_1_) as red fluorescent from cycling (S-G_2_-M) cells as GFP+ or double-positive (Fig. 1o). Scoring of red *versus* green and yellow fluorescent cells in xenografted MCF-7:FUCCI-bearing mice showed that 38.6% of cells in primary tumors were in G_0_/G_1_ phase but the value was 68.2% (*P<0.05*) in matched lung lesions (Fig. 1p,q). Immunofluorescence (IF) labeling for p27, a marker of dormancy (Bragado et al., 2013) with human-specific anti-CK8 and Ki67, revealed p27-positive and Ki67-negative human DTCs in lung sections from MCF-7 bearing mice (Fig. 1r). Hence, the majority of ER+ DTCs are arrested in the G_0_/G_1_ phase of the cell cycle, which is characteristic of dormant cells (Park and Nam, 2020).

### ER+ DTCs display organ-specific features

With the unprecedented opportunity to study dormant ER+ DTCs in vivo, we sought to visualize them within the tissue context using optically-cleared lungs and brains from NOD.Cg-*Prkdc^scid^ Il2rg^tm1Wjl^* Tg(CAG-EGFP)1Osb/SzJ (*NSG-EGFP*) mice together with the mammary glands bearing MCF-7:*RFP+* intraductal xenografts at an advanced stage (5 months) (Extended Data Fig. 2a). Fluorescence stereomicroscopy of individual lung lobes showed dispersed RFP+ foci (Fig. 2a). Confocal imaging revealed RFP+ cylindrical structures of 50-100 μm in length and 10-20 μm in diameter in the lung parenchyma (Fig. 2a,b). T47D and METS15 lung micro-metastases had similar morphologies (Extended Data Fig. 2b). Brain micro-metastases were detected in the meninges (Fig. 2c,d) and confocal imaging of anti-CK8 labeled sections identified round and dispersed tumor cells larger than the surrounding murine cells (Fig. 2d and Extended Data Fig. 2a). In brains with >10^7^ p/sec/cm^2^/sr bioluminescence signal, dispersed CK8+ cell clusters of 100-200 μm in diameter were revealed by 3D reconstruction of Z-stacks (Fig. 2e). Long cellular extensions reminiscent of a specialized type of filopodia for transport of signaling molecules, called ‘cytonemes’ (Kornberg and Roy, 2014) were observed in brain DTCs (Fig. 2f) suggesting that ER+ DTCs establish themselves by different strategies at different distant organs.

**Fig. 2.**
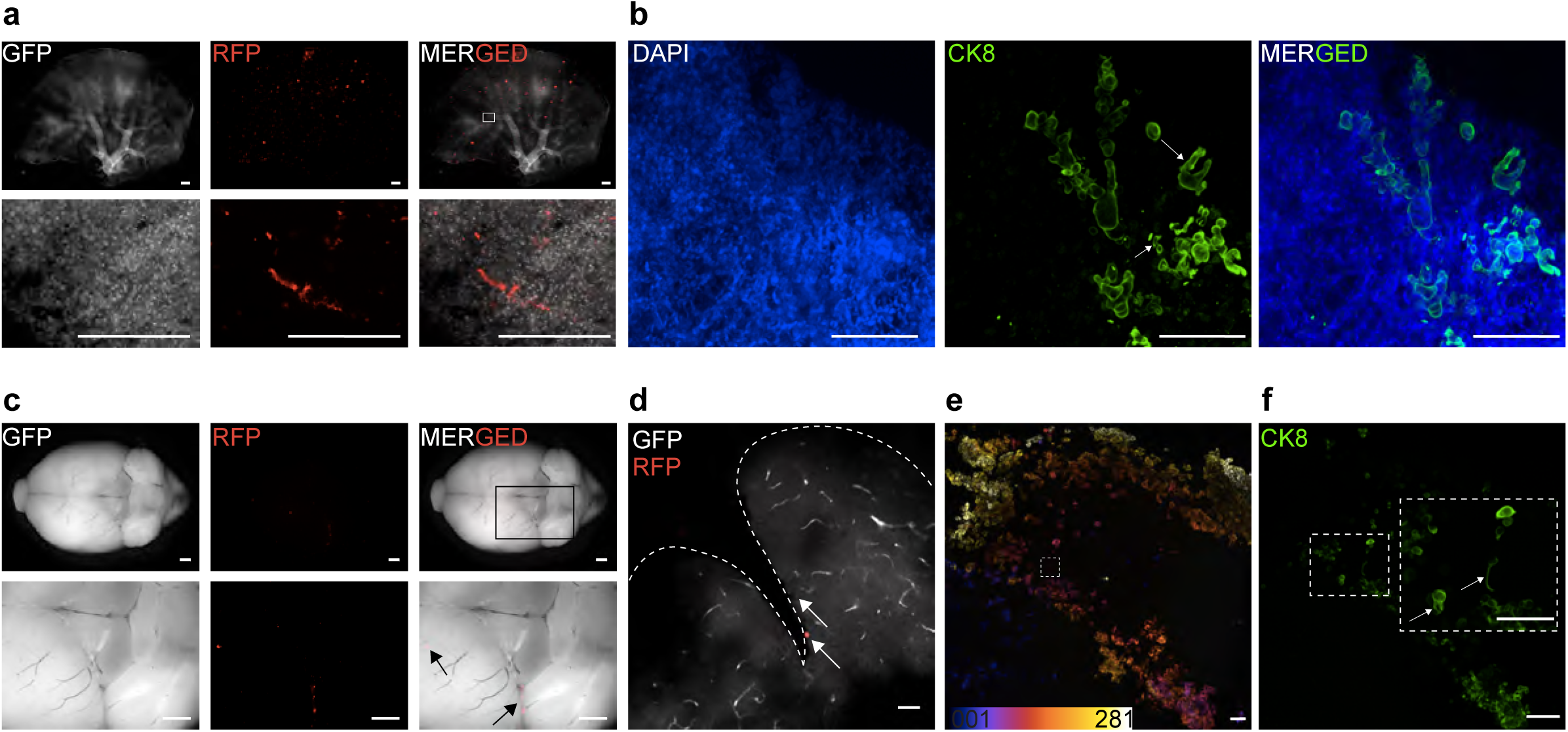
Phenotypic characterization of dormant DTCs. **a**. Fluorescence stereo micrographs of lung micro-metastases in MCF-7 intraductal xenografts-bearing mice with bioluminescence > 10^7^ p/sec/cm^2^/sr. Scale bars, 500 µm. **b**. 3D reconstruction of optically-cleared lung sections stained with CK8 and counterstained with DAPI. Scale bar, 100 µm. **c.** Fluorescence stereo micrographs of brain micro-metastases in MCF-7 intraductal xenografts-bearing mice. Scale bars, 1 mm. **d.** Fluorescence micrographs of fat-cleared brain sections from MCF-7 intraductal xenografts in NSG.GFP+ mice. Arrow points to MCF-7 cells. Scale bar, 50 µm. **e.** Depth-coded 3D reconstruction of brain micro-metastases in fat-cleared brain sections with bioluminescence > 10^7^p/sec/cm^2^/sr from MCF-7 intraductal xenograft bearing mice. Scale bar, 100 µm. Color code represents the Z-stacks. **f.** Single Z section from the hyperstack in (**f**) stained with CK8. Arrows point to cellular protrusions. Scale bar, 100 µm.

### DTCs have features of epithelial-mesenchymal plasticity (EMP)

Individual lung DTCs appeared mesenchymal (Fig. 3a) inciting us to quantify the cellular aspect ratio (AR) (major axis: minor axis) (O’Connor and Gomez, 2013) of primary tumor cells and matched lung DTCs. This showed that at the primary site, 10-15% of cells had an AR>1.7 associated with mesenchymal cells (Fig. 3b) whereas in the lungs this number increased to 40-50% (Fig. 3c).

**Fig. 3.**
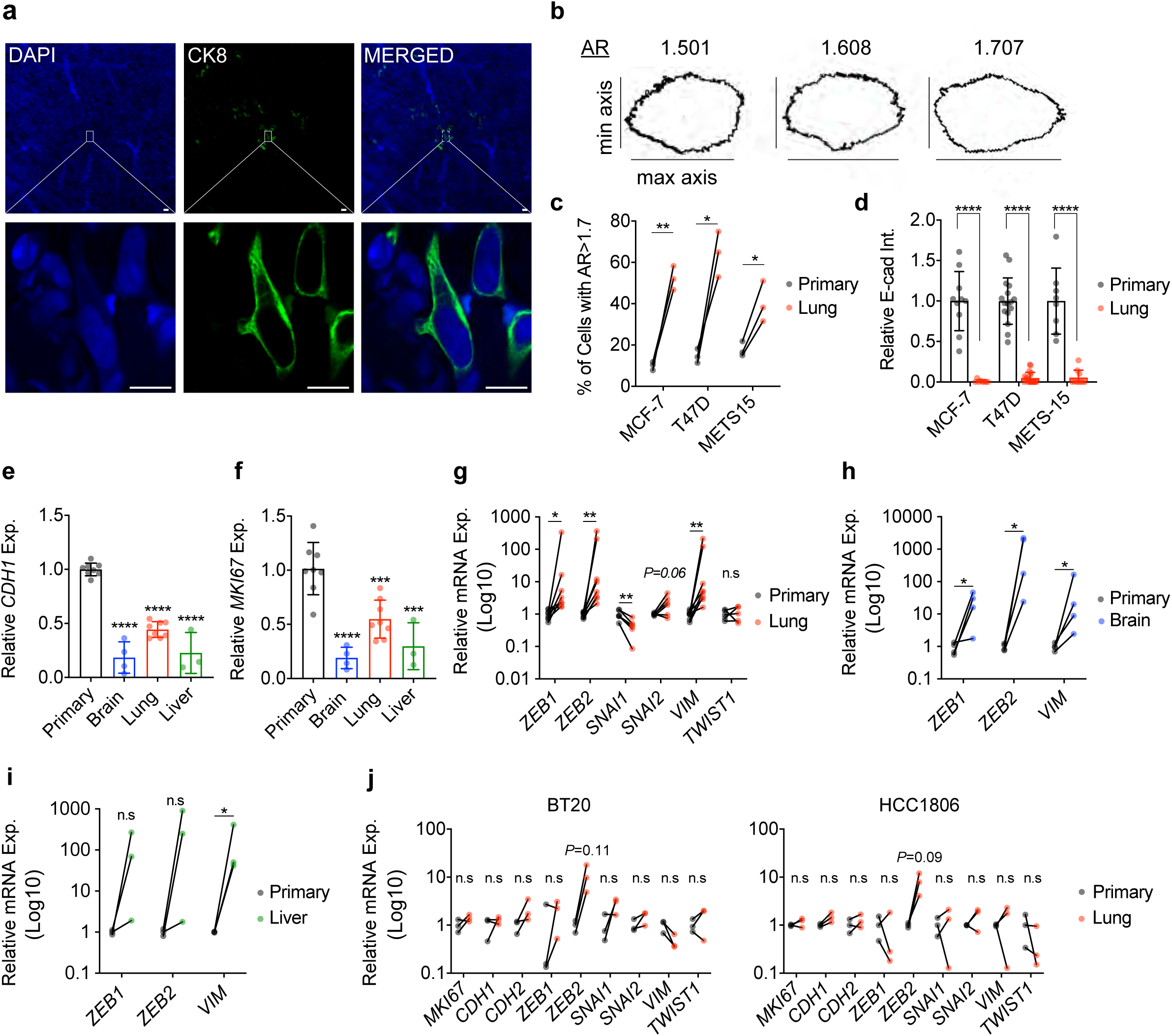
Dormant DTCs from ER+ intraductal xenografts have an EMP signature. **a.** CK8 staining on optically-cleared sections of METS15 tumors cells in host’s lungs. Scale bar, 10 µm. **b.** Representative masks representing different aspect-ratio (AR) with respect to the morphology **c.** Percentage of cells with AR > 1.7, representing mesenchymal-like cells in matched primary and lung metastatic cells from ER+ intraductal xenografts-bearing mice. Data represent mean ± SD from 3 mice. Paired Student’s t-test; **** represents p<0.0001. **d**. Relative E-cad intensity (Int.) in matched primary and lung metastatic cells from MCF-7, T47D, and METS15 intraductal xenografts-bearing mice. E-cad intensity was quantified from IHC sections of matched primary and lung metastatic cells using the corrected total cell fluorescence formula. Data represent mean ± SD from 3 mice. **e.** Relative *CDH1* and **f**. *MKI67* mRNA expression (Exp.) levels in MCF-7 cells of the primary tumor (8 mice), or in the lung (n=8), brain (n=4), and liver (n=3) DTCs. Data represent mean ± SD. One-way ANOVA. **g**. Relative mRNA expression levels in matched primary and lung DTCs (n=8), normalized to the geometric mean of *GAPDH* and *HPRT*. Wilcoxon test. **h.** Relative mRNA expression levels in matched primary and brain (**f**) n=4 and **i.** liver DTCs n=3. Wilcoxon test. **j**. Relative mRNA expression levels in matched BT20 and HCC1806 primary and lung metastases from 3 mice. Wilcoxon test. *, **, ***, ****, and n.s. represent p<0.05, 0.01, 0.001, 0.0001, and not significant, respectively.

To address whether DTCs have mesenchymal features, we analyzed E-cadherin (E-cad) protein levels. By IF, E-cad expression was readily detected in the primary tumors formed by MCF-7, T47D, and METS15 cells but reduced to < 10% in matched lung DTCs (Fig. 3d and Extended Data Fig. 3a-c). We then micro-dissected fluorescent lesions in the brain, lungs, and liver, isolated RNA and compared mRNA expression to that of the respective primary tumors using human-specific primers. Real time PCR (RT-PCR) analysis showed *MKI67* transcript levels were 80%, 45%, and 70% lower in the metastatic lesions than in the primary tumor (Fig. 3f). *CDH1* transcript levels in brain, lung, and liver lesions, were down-modulated to 81%, 56%, and 78% compared to the primary tumor (Fig. 3e). Transcript levels of the EMT transcription factors, *ZEB1* and *ZEB2*, and the mesenchymal intermediate filament *VIM* were 46-, 77-, and 46-fold higher, respectively, in lung lesions compared to primary tumor (Fig. 3g). *SNAI2* and *TWIST1* mRNA expression levels were not significantly increased, while *SNAI1* expression was down-modulated 2.3-fold (*P*<0.05) (Fig. 3g). *ZEB1, ZEB2*, and *VIM* transcript levels were also up-regulated in brain DTCs (*P<0.05*) (Fig. 3h). In the 3 liver lesions we isolated, only *VIM* transcripts were significantly up-regulated (*P*<0.05*). ZEB1 and ZEB2* transcript levels showed an increasing trend (*P*<0.12 and *P*<0.25, respectively) (Fig. 3i). Similarly, *MKI67* and *CDH1* transcript levels were 80% and 57% lower in lung micro-metastases from the ER+ PDX METS15, while *ZEB1, ZEB2,* and *VIM* transcript levels were increased 3.2-, 14.6-, and 67.9-fold, respectively (*P*<0.01, *P*<0.01, and *P=0.052*) (Extended Data Fig. 3j). *CDH2, SNAI1, SNAI2*, and *TWIST1* transcripts were not detected in either site. In the TN BT20 and HCC1806 cells, *MKI67, CDH1, CDH2*, *ZEB1*, *SNAI1*, *SNAI2*, *VIM*, and *TWIST1* transcript levels were comparable at matched primary site and the lungs (Fig. 3j) *ZEB2* transcript levels showed a trend to increase around 10-fold however that failed to reach significance (Fig. 3j). Thus, mesenchymal morphology and increase in expression of multiple EMP markers are specifically observed in ER+ DTCs.

### Role of EMP in ER+ tumor progression

Evidence that tumor cells can assume different states of EMP that are important for tumor progression has accumulated in different models (Aiello et al., 2018; Pastushenko et al., 2018; Rios et al., 2019; Simeonov et al., 2021). To assess whether EMP occurs during the transition from *in situ*to invasive disease in the ER+ MCF7 MIND model, we measured the expression levels of *CDH1* and EMT inducing transcription factors (EMT-TFs) at 1 month after injection when primary tumors are *in situ*and 5 months later when they have become invasive (Fig. 4a). *CDH1*, *ZEB1*, and *VIM* transcript levels did not change (Fig. 4b), *ZEB2* levels decreased by 50% (*P<0.05)*, and *TWIST1* and *SNAI2* levels increased 15.6 and 8.6-fold (*P<0.01* and *P<0.05*), respectively (Fig. 4b) at the invasive stage. IF showed that MCF-7 cells also retained E-cad protein expression at the invasive stage (Fig. 4c).

**Fig. 4.**
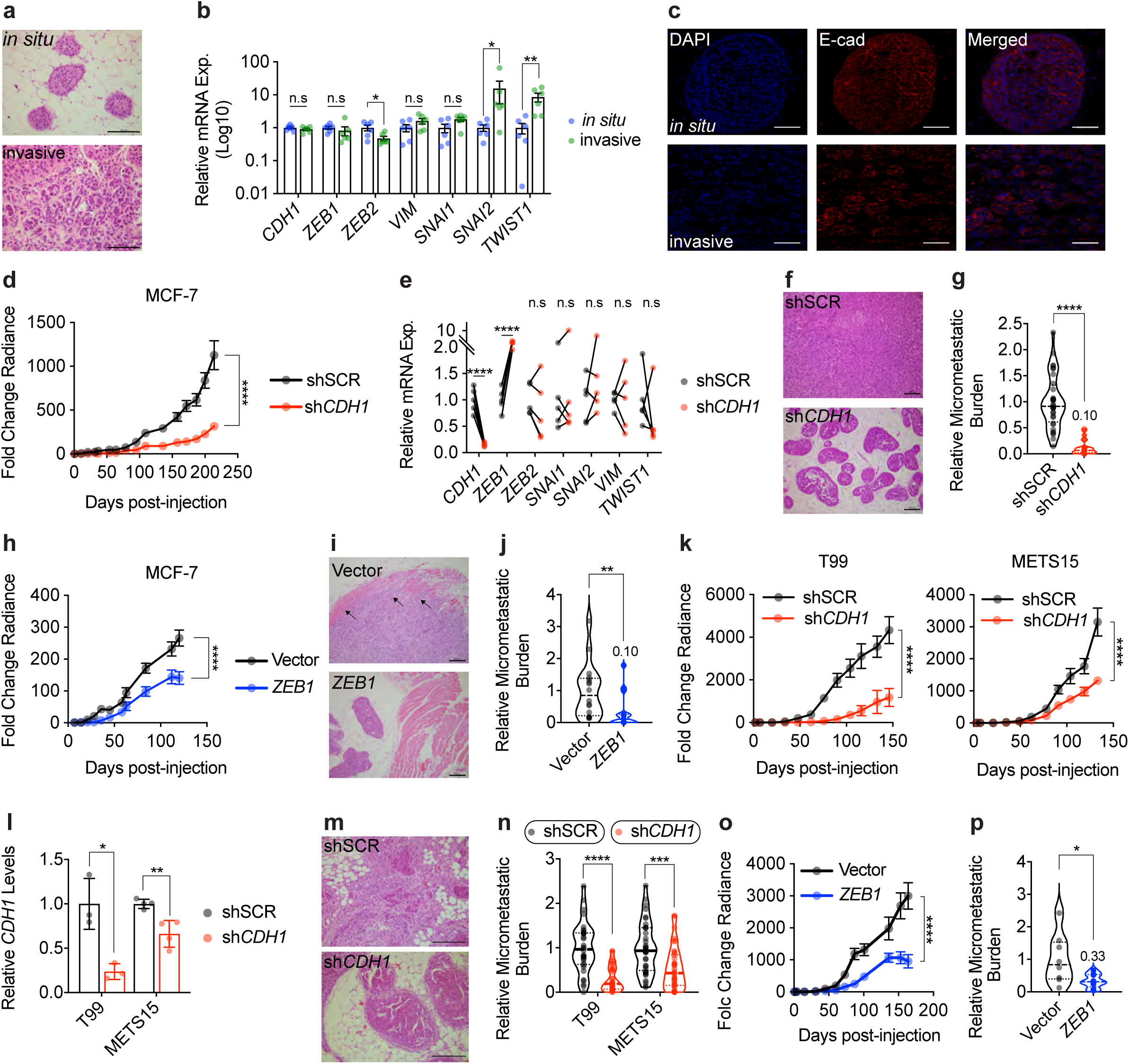
Role of EMP in ER+ tumor progression. **a.** Representative H&E micrographs of *in situ*(1 month after injection) and invasive (5-6 months after injection) MCF-7 intraductal xenografts, scale bars, 100 µm. **b.** Relative mRNA expression levels of the selected genes in *in situ*and invasive MCF-7 intraductal xenografts, normalized to the geometric mean of *GAPDH* and *TBP*. Data represent mean ± SD from 6 glands in 3 mice, non-parametric Mann Whitney test. **c**. Representative IF micrographs of E-cad in *in situ*and invasive MCF-7 intraductal xenografts. Nuclei were counterstained with DAPI, scale bars, 50 µm. **d.** Graph showing the fold change bioluminescence over time of MCF-7 shSCR and sh*CDH1* in intraductal xenografts, data represent mean ± SEM of 20 and 19 xenograft glands, respectively, 5 mice for each group. Two-way ANOVA, multiple comparisons. **e.** Relative mRNA expression levels of selected genes in 5 contralateral shSCR and sh*CDH1* xenograft glands, normalized to the geometric mean of *GAPDH* and *TBP*. Paired t-test. **f.** Representative H&E micrograph from shSCR and sh*CDH1* xenograft glands. Scale bars, 200 µm. **g.** Relative micro-metastatic burden in 7 sh*SCR* and 8 sh*CDH1* tumor-bearing mice. Each dot represents a single organ, the dashed line represents the median and dotted lines the lower and upper quartiles. Unpaired Student’s t-test. **h.** Graph showing the fold change bioluminescence over time of Vector and *ZEB1*-overexpressing MCF-7 cells in intraductal xenografts, data represent mean ± SEM of 16 xenograft glands from 4 mice each. Two-way ANOVA, multiple comparisons. **i.** Representative H&E micrographs from thoracic mammary glands bearing either vector or *ZEB1*-overexpressing MCF-7 cells from 3 mice, each. Arrows point to invaded muscle tissue in thoracic mammary glands. Scale bar, 200 µm. **j.** Violin plot of the relative micro-metastatic burden from vector and *ZEB1*-overexpressing MCF-7 intraductal xenografts-bearing mice, respectively. The dashed line represents the median and dotted lines indicate the lower and upper quartiles. Unpaired Student’s t-test. **k.** Growth curve of intraductal T99 and METS15 shSCR or sh*CDH1*. For T99, data represent mean ± SEM of 16 xenograft glands for each shSCR and sh*CDH1* groups, 5 mice for each cohort, and 16 and 20 xenograft glands for METS15, respectively, 5 mice for each cohort. Two-way ANOVA, multiple comparisons. **l.** Relative *CDH1* expression in 3 T99 shSCR and sh*CDH1* intraductal xenografts and 4 METS15 shSCR and sh*CDH1*. Data represent mean ± SD. Unpaired Student’s t-test. **m.** Representative H&E micrographs of T99 shSCR and sh*CDH1* xenograft glands. Scale bar, 200 µm. **n.** Violin plot of the relative micro-metastatic burden in T99 and METS15 intraductal xenografts-bearing mice. The dashed line represents the median and dotted lines the lower and upper quartiles. Each dot represents a single organ. Unpaired Student’s t-test. **o.** Growth curve of vector control and *ZEB1*-overexpressing T99 intraductal xenografts. Data represent mean ± SEM of 15 and 18 xenograft glands, respectively. Two-way ANOVA, multiple comparisons. **p.** Violin plot of the relative micro-metastatic burden in the lungs and bones from 4 Vector and 5 *ZEB1*-overexpressing T99 xenografts-bearing mice. The dashed line represents the median and dotted lines the lower and upper quartiles. Unpaired Student’s t-test. *, **, ***, ***, and n.s represent *P*<0.05, 0.01, 0.001, 0.0001, and not significant, respectively.

To test functionally whether EMP favors the metastatic spread of ER+ BC cells, we induced EMP by two complementary approaches; in the first, we down-modulated *CDH1* expression, and in the second, we overexpressed *ZEB1* in MCF7 cells. Given the heterogeneity of the lentivirally transduced cell population different states of EMP likely result in different cells by this approach. Morphological assessment and expression analysis of EMT-TFs showed that both MCF-7:sh*CDH1* and MCF-7:*ZEB* cells acquired EMT features according to the recent guidelines (Yang et al., 2020) (Extended Data Fig. 4a-h).

When MCF-7:sh*CDH1* cells were grafted intraductally, they grew approximately 3.5-fold less than the MCF-7:shSCR controls (Fig. 4d) and mammary gland weight at end-point was reduced by 73% (P<0.0001) (Extended Data Fig. 4i). At sacrifice, E-cad expression in the xenografts was reduced by 75% as determined by Western blot and immunohistochemistry IHC (Extended Data Fig. 4j,k). Ki67 and pHH3 indices were decreased by 30% and 75% in MCF-7:sh*CDH1* compared to MCF-7:shSCR grafts, respectively (Extended Data Fig. 4l). MCF-7:sh*CDH1* cells similarly maintained a mesenchymal phenotype and a 3-fold increase in *ZEB1* (Fig. 4e).

Analysis of hematoxylin and eosin (H&E) stained sections showed that MCF-7:shSCR cells invaded the stroma whereas the MCF-7:sh*CDH1* cells remained mostly *in situ*(Fig. 4f). The micro-metastatic load of mice engrafted with MCF-7:sh*CDH1* cells was around 10% of that of mice with MCF-7:sh*SCR* cells (Fig. 4g and Extended Data Fig. 4m). To exclude the possibility that the decreased metastatic burden merely reflects decreased primary tumor growth, we performed a paired analysis of the micro-metastatic burden of mice with similar tumor burden at endpoint. In these mice, the micro-metastatic load in the brain, lungs, liver, and bones was reduced by 90% (Extended Data Fig. 4n).

To exclude potential systemic effects of the genetically different tumor cells on the host organism, we contralaterally grafted MCF-7:sh*SCR:GFP* and MCF-7:sh*CDH1:RFP* cells (Extended Data Fig. 4o). Comparison within the same host showed that the signal emanating from glands engrafted with MCF-7:sh*CDH1:RFP* cells at endpoint was 6- and 2-fold lower than that from contra lateral glands as determined by *in vivo* imaging and *ex vivo* fluorescence, respectively (Extended Data Fig. 4p). The number of sh*CDH1:* RFP+ lung foci was 8-fold lower than the number of GFP+ loci and the total RFP+ area was 15-fold smaller than the respective GFP+ area (Extended Data Fig. 4q, r).

Similarly, MCF-7:*ZEB1* xenografts grew 2-fold less than MCF-7:*Ctrl* xenografts (Fig. 4h) and the corresponding xenografted glands weighed 1.5-fold less than their controls (*P<0.05*) (Extended Data Fig. 4s) when host mice were sacrificed at 4 months. Histological examination revealed that control cells invaded the muscle tissue adjacent to the thoracic mammary gland, whereas *ZEB1*-overexpressing cells remained *in situ*with occasional microinvasion foci (Fig. 4i). In addition, *ZEB1* overexpression reduced the overall micro-metastatic burden by 90% (Fig. 4j) to a varying extent in different organs (Extended Data Fig. 4t).

Similarly, down-modulation of E-cad expression in the ER+ PDXs, T99 and METS15 reduced *in vivo* growth (Fig. 4k) and mammary gland weight at end-point (Extended Data Fig. 4u). RT-PCR analysis revealed a 77% and 33% down-modulation of *CDH1* levels in the glands xenografted with T99 and METS15 cells, respectively (Fig. 4l). Furthermore, E-cad down-modulation reduced tumor invasion (Fig. 4m) and micro-metastatic burden (Fig. 4n and Extended Data Fig. 4v) of both ER+ PDXs. Together these data indicate that E-cad is required for metastasis in ER+ BC.

Ectopic *ZEB1* expression decreased T99 intraductal growth 3-fold, (Fig. 4o) and reduced the weight of xenografted glands (Extended Data Fig. 4w) and the micro-metastatic load (Fig. 4p) both in the lungs and bones (Extended Data Fig. 4x). Therefore, EMP *per se*, induced by *CDH1* down-modulation or *ZEB1* overexpression, does not favor tumor progression nor metastasis of ER+ BC cells. These findings suggest that DTCs are indeed plastic and the mesenchymal phenotype may be induced after cells leave the primary tumor, likely at the distant sites.

### E-cadherin restoration awakens DTCs from dormancy

Our finding that dormant DTCs are in EMP states prompted us to assess whether a return to an epithelial state may reactivate proliferation. To this aim, we dissociated mammary glands and lungs from MCF7:*RFP* in *NSG-EGFP* mice to single cells, plated them in 2D and applied drug selection to avoid overgrowth of the abundant mouse cells. MCF-7:*RFP+* cells derived from the primary tumor proliferated within a few days and were confluent by 1-2 weeks of *in vitro* culture (Extended Data Fig. 5b). The lung-derived DTCs gradually resumed proliferation after 2-3 months in culture, and formed epithelial islets (Extended Data Fig. 5a). *CDH1* transcript levels were restored and expression of EMT-TFs decreased (Extended Data Fig. 5c,d) which is consistent with the hypothesis that reacquisition of an epithelial state enables cell proliferation.

To address whether restoration of an epithelial state is sufficient to awaken dormant DTCs in vivo, we overexpressed E-cad in MCF-7 cells (Extended Data Fig. 5e). In vivo bioluminescence 1 day after intraductal engraftment was 2-fold higher in glands engrafted with MCF-7:*CDH1* cells compared to control grafts suggesting that E-cad favors tumor cell survival and/or engraftment in the mouse milk ducts (Fig. 5a). Six months after injection, however, bioluminescence in MCF-7:*CDH1* grafts was 2-fold lower than that of the controls (Fig. 5b). Similar to the 10-fold increase initially observed *in vitro* (Fig. 5c and Extended Data Fig. 5e), an 8-fold *CDH1* transcript overexpression was observed. *Ex vivo* bioluminescence in the lungs from mice with MCF-7:*CDH1* xenografts was 10% that of MCF-7:control xenograft bearing mice (P<0.05) (Fig. 5d). Yet, in 1 of the 4 mice engrafted with MCF-7:*CDH1*, a lung lesion > 500 µm in diameter was detected (Fig. 5e), when such a large lesion was never observed in more than 50 mice engrafted with parental MCF-7 cells. This finding suggested the possibility that E-cad expression may have enabled an exit from dormancy.

**Fig. 5.**
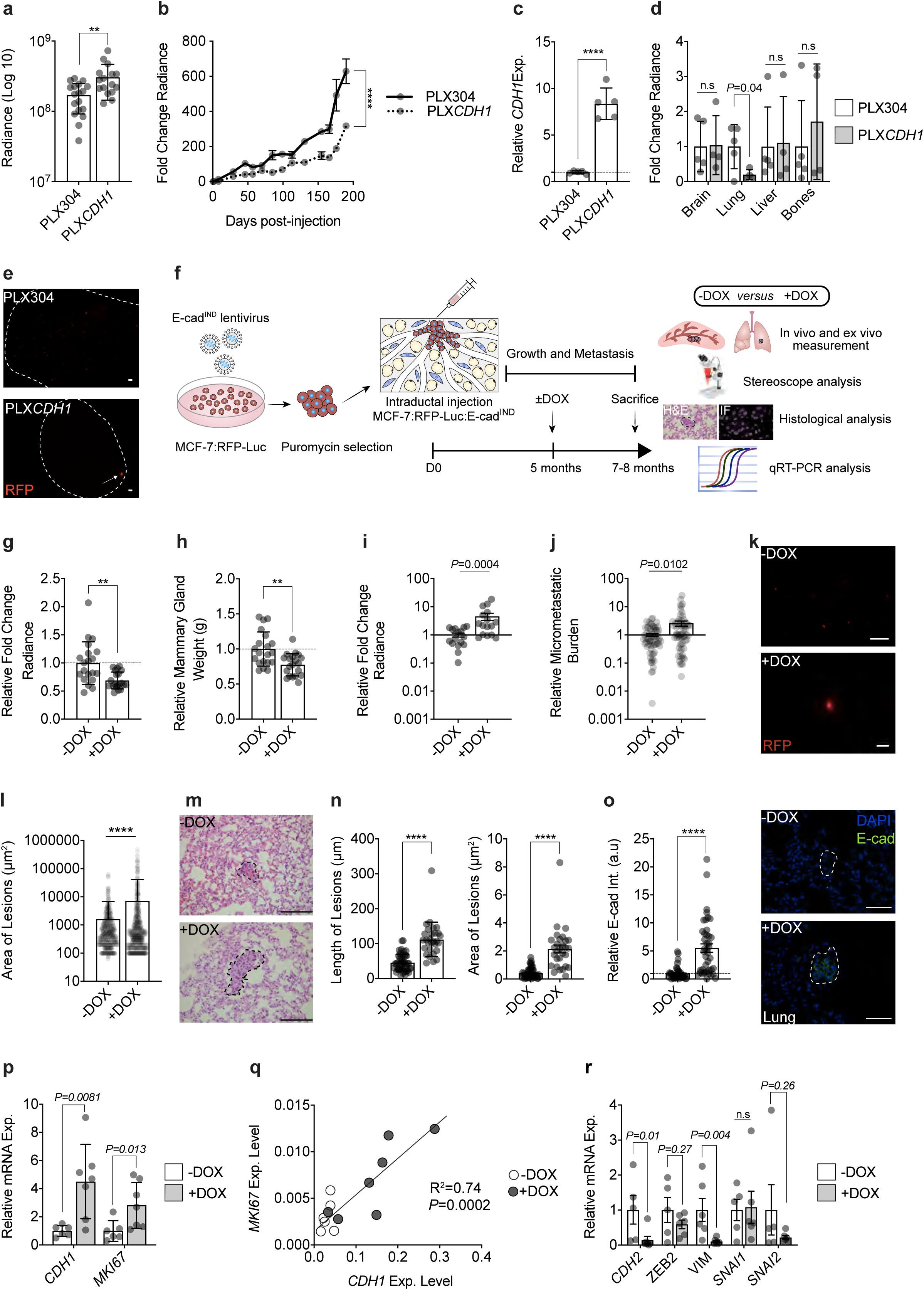
Role of E-cadherin restoration in tumor cell awakening. **a**. Bioluminescence values on Day 1 after intraductal injection of control (PLX304) and *CDH1*-overexpressing (PLX*CDH1*). Data represent mean ±SD of 17 and 16 xenografted glands, respectively in 5 and 4 mice, respectively. Unpaired Student’s t-test. **b**. Fold change bioluminescence measurements of PLX304 and PLX*CDH1*-MCF-7 cells in 5 and 4 mice, respectively. Data represent mean ± SEM of 17 control and 16 *CDH1*-overexpressing intraductal xenografts from 5 and 4 mice, respectively. **c**. Relative *CDH1* levels in 5 tumors from PLX304 and PLX*CDH1*-bearing mice. Data represent mean ± SD. Unpaired Student’s t-test. **d**. Fold change bioluminescence of ex vivo bioluminescence in 5 PLX304 and 4 PLX*CDH1*-bearing mice. Data represent mean ± SD. Multiple t-tests. **e**. Fluorescence stereo microscope micrographs of lung lesions in PLX304 (top) and PLX*CDH1* (bottom). 1 out of the 5 analyzed mice had a lesion >500 µm shown over here. Scale bar, 500 µm. **f**. Experimental scheme of the intraductal injection of MCF-7:RFP-luc:E-cad^IND^. **g**. Relative bioluminescence and **h**. weight of 20 xenografted glands from 9 (-DOX) and 9 (+DOX) mice. Data represent mean ± SD. Unpaired Student’s t-test. **i**. Fold change lung bioluminescence in -DOX and +DOX from 16 mice. Data represent mean ± SEM. Non-parametric Mann-Whitney Test. **j**. Relative micro-metastatic burden in 16 -DOX and +DOX mice. Data represent mean ±SEM and each organ represents a single organ. Non-parametric Mann-Whitney Test. **k.** Fluorescence stereo micrographs of lung lesions in -DOX (top panel) and +DOX (bottom panel) treated mice. Scale bars, 500 µm. **l.** Quantification of the area of lesions in lungs from mice treated with (-DOX) or (+DOX) n=9 by fluorescence stereomicroscopy. Data represent mean ± SD. Unpaired Student’s t-test. **m.** Representative H&E lung micrographs from 3 mice for -DOX and +DOX. Scale bar, 100 µm. **n.** Quantification of the length (left) and area of lesions (right) in H&E lung sections from 3 (- DOX) and 3 (+DOX) mice. Data represent mean ± SD. Unpaired Student’s t-test. **o.** Relative E-cad intensity (Int.) in lung lesions and representative IF of E-cad in lung lesions from 3 - DOX and 3 +DOX mice. Nuclei were counterstained with DAPI. Scale bar, 50 µm. Data represent mean ± SEM. Unpaired Student’s t-test. **p**. Relative mRNA levels of *CDH1* and *MKI67* normalized to *GAPDH* in lungs from 6 -DOX and 7 +DOX mice. Data represent mean ± SD. Non-parametric Mann Whitney test. **q.** Plot showing correlation between *MKI67* and *CDH1* expression levels in 6 -DOX and 7 +DOX mice. **r.** Relative mRNA expression levels of EMT-TFs in flash-frozen lungs from 5 -DOX and 6 +DOX mice, normalized to *GAPDH*. Data represent mean ± SD. Non-parametric Mann Whitney test. *, **, ***, ***, and n.s represent p<0.05, 0.01, 0.001, 0.0001, and not significant, respectively.

To circumvent the confounding effects of E-cad overexpression on engraftment and *in vivo* growth of MCF-7 cells at the primary tumor site, we used a doxycycline (DOX) inducible E-cad vector (E-cad^IND^) (Extended Data Fig. 5f). Five months after engraftment of MCF-7:*E-cad^IND^* cells, when DTCs are readily detected in the lungs, a subgroup of mice was switched to DOX-containing chow for 2-3 months (Fig. 5f). Bioluminescence and weight of mammary glands at sacrifice were significantly reduced in the DOX-induced group (*P<0.01*) (Fig. 5g,h), but lung bioluminescence increased by 4.5-fold (Fig. 5i) and the overall bioluminescence signal in distant organs by 4-fold (Fig. 5j). To assess whether the increased lung bioluminescence was due to increased seeding and/or an increase in size of individual lesions, we measured the micro-metastases by fluorescence stereo microscopy and histological analysis. *CDH1* induction resulted in a 4.5-fold increase in the overall fluorescent area (Fig. 5k,l). Similarly, quantitative image analysis of lung lesions in H&E-stained sections revealed a 5.3 and 2.5-fold increase in their area and length, respectively (Fig. 5m,n). Quantification of E-cad intensity of lung micro-metastases from control and +DOX mice showed a 5.5-fold increase in protein expression in the lung micro-metastases upon DOX administration (Fig. 5o). Semi-quantitative RT-PCR on lungs from control and +DOX groups confirmed induction of *CDH1* and revealed a 2.8-fold increase in *MKI67* expression levels (Fig. 5p). *CDH1* expression correlated positively with *MKI67* expression (P<0.001) (Fig. 5q), reinforcing the notion that E-cad restoration drives cell cycle entry and proliferation. Finally, we asked whether the increase in proliferation in DOX-treated mice involved loss of mesenchymal phenotype. We noted a significant decrease in *CDH2* (*P*=0.01) and *VIM* (*P*=0.004) transcript levels upon *CDH1* restoration (Fig. 5r). *ZEB2* and *SNAI2* transcript levels showed a decreased trend while *SNAI1* levels were not altered (Fig. 5r). *ZEB1* was not reliably detected by the semi-quantitative RT-PCR approach (cycle threshold >37). Thus, E-cad overexpression with resulting MET is sufficient to drive DTCs out of dormancy. Our data indicate that EMP and not a single epithelial or mesenchymal cell state is responsible for distant metastasis, tumor cell dormany and latent disease and targeting of this plastic process provides a new therapeutic angle for tertiary prevention in ER+ BC (Fig. 6).

**Fig. 6.**
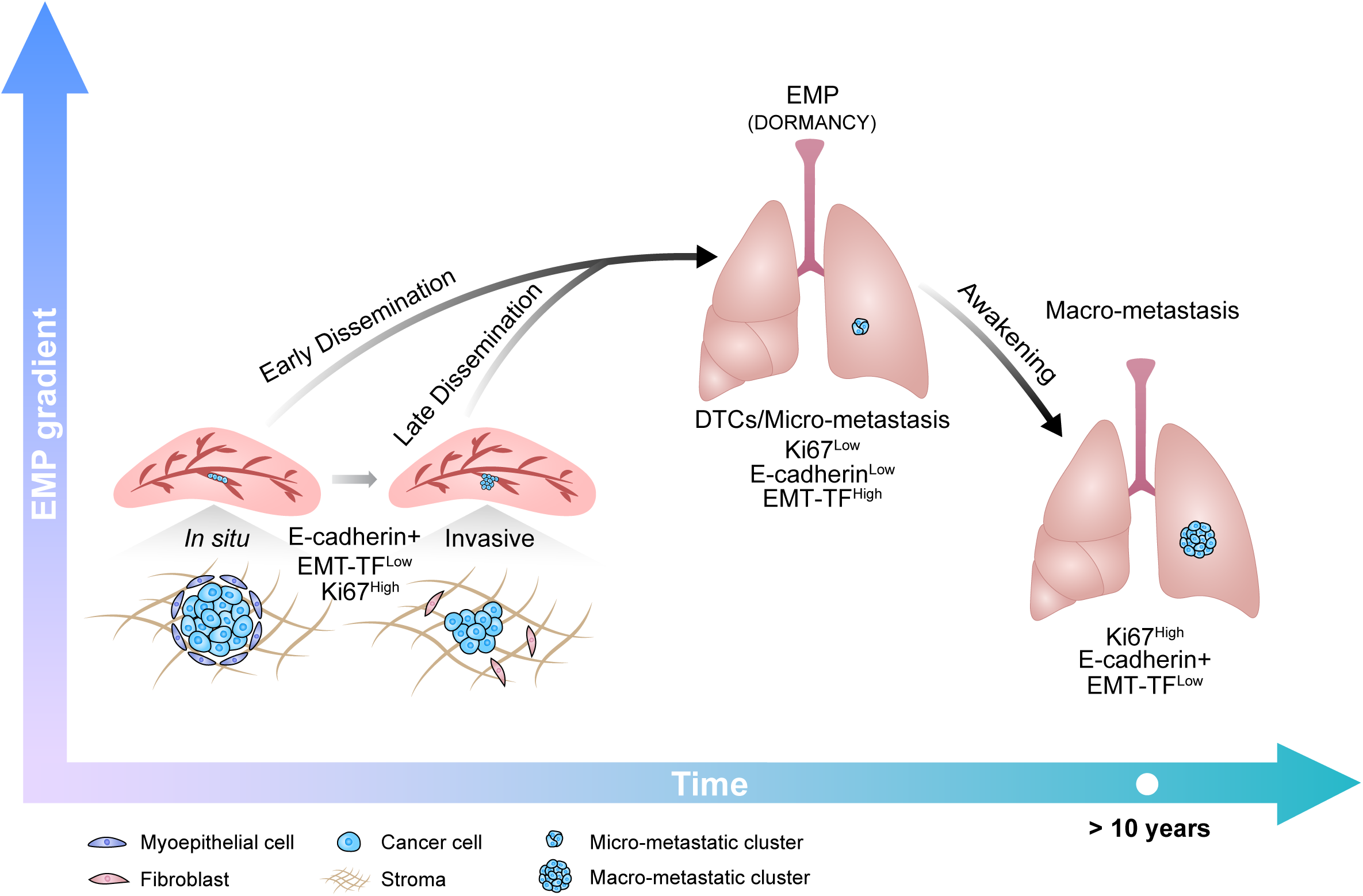
Working Model for ER+ BC progression. ER+ tumor cells progress from the *in situ* to the invasive stage in an epithelial state characterized by high-level expression E-cadherin and low expression of EMT-TFs. At distant sites, DTCs show EMP features and enter dormancy. Awakening from dormancy involves restoration of E-cadherin and suppression of the EMT-TFs. Drugs promoting EMP or inhibiting the reacquisition of an epithelial state are warranted to prevent disease recurrence. EMT-TFs: Epithelial-MesenchymalTr ansition-activating transcription factors; EMP: Epithelial-Mesenchymal Plasticity; DTCs: Disseminated Tumor Cells

## Discussion

Early tumor cell dissemination and dormancy in ER+ BC (Buell et al., 2004; Matser et al., 2018; Sänger et al., 2011) represent important clinical challenges. Our understanding of the mechanisms controlling these phenomena is very limited because of the lack of adequate preclinical models to study them. Here, we establish with intraductal BC xenograft models that TN DTCs form macro-metastases whereas specifically ER+ DTCs enter dormancy. As the engrafted cells, cell line or patient-derived can be readily genetically manipulated, our approach opens new opportunities to study the entire metastatic process mechanistically.

The high proliferative indices of TN versus ER+ DTCs we demonstrate *in vivo*, provide an explanation for the success of chemotherapy in TN BC patients, which can kill proliferating DTCs and incipient metastases. In contrast, ER+ BC patients generally respond poorly with some benefit observed in patients with the more highly proliferative luminal B subtype (Criscitiello et al., 2014). Our finding that the proportion of tumor cells residing in the G_0_/G_1_ phase of the cell cycle is far higher in ER+ DTCs than in the primary tumor, albeit not 100% is in line with their being a gradient of different EMP states and different proliferative states within a given tumor. In the clinical settings, the definition of luminal B versus A subtype is based on Ki67 indices and the cut-off has been much discussed (Urruticoechea et al., 2005). This may reflect a similar spectrum of different states of cellular plasticity and proliferative potentials. It is conceivable that the more proliferative subset of DTCs in patietns with luminal B tumors is abrogated by the chemotherapy resulting in improved survival rates.

Our finding that the epithelial differentiation state reflected by E-cad expression is critical for recurrences in ER+ BC adds to previous evidence that it is important to tumor progression in different models (Dong et al., 2007; Hugo et al., 2017; Kleer et al., 2001; Kowalski et al., 2003; Padmanaban et al., 2019). It has been postulated that EMP is induced at the distant sites based on observations with a syngeneic mouse mammary tumor cell line model, which grows in the lungs following intravenous injection (Montagner et al., 2020). Recent demonstrations that E-cad expression persists in CTCs (Koch et al., 2020; Na et al., 2020) and increases their survival (Padmanaban et al., 2019) further point in the direction of a late EMP. Our observations are compatible with such a scenario but we cannot exclude the existence of subpopulations of cells with EMP at the primary site or as circulating tumor cells. Interestingly, we observed that in the primary tumor both E-cad upregulation and down modulation reduced tumor growth. This suggests that E-cad levels need to be tightly regulated and highlights the importance of of doing functional studies in the context of of the DTCs rather than on primary tumors.

The upregulation of *SNAI2* and *TWIST1* transcripts in the invasive versus *in situ* disease, is in line with previous findings that the fat pad microenvironment induces these factors as part of a basal differentiation program triggered by TGFβ-signaling (Sflomos et al., 2016). As such, their elevated expression is likely a consequence of invasion and not causally related to it. In line with cell type-specific EMP states (Pastushenko et al., 2018), our findings suggest that in ER+ human BC cells the critical EMT-TFs are *ZEB1/ZEB2* whereas *SNAI2* and *TWIST1,* critical for EMP in other tissues, control a distinct cellular differentiation program that is linked to basal features. While we find *ZEB1/ZEB2* transcripts increased in all organs sites further experiments are needed to determine organ-specific aspects of the EMP that may ultimately also be exploited therapeutically.

The E-cad down-modulation observed in dormant DTCs is most likely due to the ability of EMT-TFs to directly bind the *CDH1* promoter and repress its transcription (Batlle et al., 2000; Comijn et al., 2001). Of note, EMT-TFs, ZEB1/ZEB2, SNAIL, and TWIST, as well as EMT-inducing factors, such as TGFβ, were all shown to halt the cell cycle by decreasing Cyclin D protein levels and increasing the phosphorylation of Rb (Lovisa et al., 2015; Mejlvang et al., 2007; Vega et al., 2004; Yang et al., 2006). Interestingly, ZEB1 can repress ER-α expression by inducing the hypermethylation of its promoter (Zhang et al., 2017) and might be responsible for the decrease in ER expression. *ZEB1* expression can also be activated by hormones namely, estrogens (Chamberlain and Sanders, 1999), progesterone (Richer et al., 2002), and androgens (Anose and Sanders, 2011), pointing to a tight connection between Zeb1 and HR signaling. It is tempting to speculate that the reduced recurrence and breast cancer-related mortality in patients who receive tamoxifen for 10 years compared to only 5 years (Davies et al., 2013; Gray et al., 2013) is due to a sustained EMP-induced dormant state. Indeed, evidence has accumulated that endocrine therapy induces EMT signatures in ER+ BC (Creighton et al., 2009; Kastrati et al., 2020; Sflomos et al., 2016).

A limitation of the intraductal xenograft models used in the present study is the absence of an intact immune system. NSG mice lack functional B and T cells; while some innate immune cells are present their function might be altered or incomplete in the absence of the adaptive response. This study has revealed the reacquisition of an epithelial state as critical for awakening from dormancy but it may be a multifactorial process, involving multiple cell types (Montagner et al., 2020; Nobre et al., 2020; Ombrato et al., 2019; Werner-Klein et al., 2020) and secreted factors, including cytokines and chemokines. In addition, whether active immune-cell signaling may ultimately regulate the expression of *CDH1* or other targets needs to be determined. This can be addressed by restoring a functional human immune system in the host mice in the context of humanized mouse models (Kalscheuer et al., 2012).

The observation that ER+ PDXs, no matter whether established from a primary tumor (T99) or a metastatic lesion (METS15), disseminate and enter dormancy, suggests that these processes are not genetically but epigenetically controlled. The present findings suggest that promoting a more mesenchymal cell state is critical to prevent recurrence and that in case of endocrine resistant disease, E-cad protein might by targeted directly via monoclonal antibodies (Brouxhon et al., 2013). This and other hypotheses are now readily amenable to experimental testing with intraductal xenograft models.

---

**Extended Data Fig. 1. ER+ and ER- BC cells show different metastatic behavior.**

**a.** Bar graph showing take rates for TNBC cells (blue) either cell lines (BT20 and HCC1806) or PDX, T70, and ER+ BC cells (red) either cell lines (MCF-7 and T47D) or PDXs, T99 and METS15, injected intraductally calculated as % successfully-engrafted glands of total number of injected mammary glands. Data represents mean ± SD from 41 MCF-7, 36 T47D, 32 T99, 36 METS15, 18 BT20, 16 HCC1806, and 10 T70 intraductal xenografts. The vertical dashed line equals 90% engraftment rate. **b.** Fold change bioluminescence at 5 weeks of ER+ and TNBC intraductal xenografts from **Figure 1b**. Data represent mean ± SD. Blue and red dashed lines represent the average fold change bioluminescence of TNBC (=1148) ER+ (=32), respectively. **c.** Fold change of bioluminescence at endpoint for TN (5 weeks) and ER+ (5-6 months) BC cells. Data represent mean ± SD of 17, 13, 19, 16, 18, 16, and 19 tumors from 5, 5, 5, 9, 7, and 5 mice bearing MCF-7, T47D, T99, METS15, BT20, HCC1806, and T70 xenografts, respectively. Box and whiskers represent minimum and maximum values. **d.** Representative fluorescence stereo micrographs of mammary glands 1 week (top) and 2 weeks (bottom) after intraductal injection with BT20 and **e.** HCC1806. Scale bars, 1 mm. **f.** Representative H&E micrographs of BT20 and **g.** HCC1806 intraductal xenografts 1 week (top) and 2 weeks (bottom) after injection. Scale bars, 1 mm. **h**. Ex vivo bioluminescence on organs from mice bearing BT20 and HCC1806, 1 week (left) from 10 and 4 mice, respectively, and 2 weeks (right) after intraductal injection from 4 mice. **i.** Fluorescence stereo micrographs of lungs from BT20 intraductal xenografts-bearing mice 1 week (top panel) and 2 weeks (bottom panel) after intraductal injection. The arrow points to TN DTCs. Scale bars, 50 µm. **j.** Representative IF for CK8 (green) and Ki67 (red) counterstained with DAPI (blue) of matched primary and lung metastases of 3 TNBC and **k**. ER+ intraductal xenografts-bearing mice. Scale bars, 50 µm. **, ***, ***, and n.s represent *P*<0.01, 0.001, 0.0001, and not significant, respectively.

**Extended Data Fig. 2. a.** CK8 IF on optically-cleared vibratome sections of MCF-7 primary site, lung, and brain organs. Nuclei are counterstained with DAPI. Scale bar, 50 µm. **b.** Representative fluorescence stereo micrographs of lungs from mice bearing MCF-7, T47D, and METS15 intraductal xenografts. Arrows point to atypical fibroblastic cellular morphologies. Scale bar, 50 µm.

**Extended Data Fig. 3. a.** Representative immunofluorescence micrographs for CK8 (green) and E-cad (Red) counterstained with DAPI (blue) from 3 matched MCF-7, **b.** T47D and **c.** METS15 primary and lung DTCs. Scale bar, 50 µm. **d.** Relative mRNA expression levels in matched METS15 primary and lung micro-metastases (5 mice) normalized to the geometric mean of *GAPDH* and *HPRT*. Paired t-test. * and ** represent *P*<0.05, and 0.01, respectively.

**Extended Data Fig. 4. In vitro and in vivo EMP characterization. a**. Representative Western blot of total cell lysates from *in vitro* MCF-7 cells transduced with either shSCR or sh*CDH1*-encoding lentiviruses and probed for E-cad and Lamin B1 from 4 biological replicates. **b.** Phase contrast micrograph of shSCR and sh*CDH1* MCF-7 cells, scale bar 100 µm. **c.** Relative proliferation (%) of shSCR and sh*CDH1* day 3 after seeding assessed by MTT assay. Data represent mean ± SD of 4 different biological replicates. **d.** Relative mRNA expression levels of the shown genes in shSCR and sh*CDH1* MCF-7 cells. Data represent mean ± SD of 3 different biological replicates. **e.** Western blot of Zeb1, E-cad, ER, and Lamin B1 in vector and MCF-7-*ZEB1* cells. **f.** Representative phase contrast micrograph of vector and *ZEB1* MCF-7 cells, scale bar 100 µm**. g.** Fold-change absorbance of Vector and MCF-7-*ZEB1* cells over time. Data represent mean of 3 biological replicates ± SD. **h.** Relative mRNA expression levels of the shown genes in Vector and *ZEB1* MCF-7 cells, 3-4 biological replicates. **i.** Weight of glands xenografted with either shSCR or sh*CDH1* MCF-7 cells at end-point from 46 shSCR and 43 sh*CDH1* xenograft glands, total of 25 mice. Data represent mean ± SD, unpaired Student’s t-test. **j.** Relative E-cad expression levels by Western blot in 12 shSCR and sh*CDH1* xenograft glands from at least 5 mice. Student’s unpaired t-test. **k.** Representative IF of E-cad staining in shSCR and sh*CDH1* xenograft glands from contralateral mice (n=4). Nuclei are counterstained with DAPI. Scale bar, 100 µm. **l.** Percentage of Ki67 and pHH3-positive cells in contralaterally-engrafted shSCR and sh*CDH1* and their representative IFs. Scale bar, 100 µm. Each dot represents at least 100 cells per analyzed image, 4 mice. Paired Student’s t-test. **m.** Relative fold change in ex vivo organ bioluminescence in 7 shSCR and 8 sh*CDH1* tumor-bearing mice. Data represent mean ± SD, multiple t-tests. **n.** Relative fold change in ex vivo organ bioluminescence from 5 *WT* and 5 sh*CDH1* tumor-bearing mice with similar bioluminescence at end-point. Data represent mean ± SD, multiple t-tests. **o.** Scheme showing the contra lateral engraftment of green shSCR (left) and red sh*CDH1* (right) MCF-7 cells. **p.** Bar plots showing the fold change bioluminescence (left) and total fluorescence (right) of contra lateral 8 shSCR:GFP and sh*CDH1*:RFP intraductal xenografts. Data represent mean ± SD, paired t-test. **q.** Representative fluorescence stereograph of green shSCR and red sh*CDH1* cells contralaterally-engrafted. **r.** Quantification of total red and green lung foci (left) and their percentage area (right) occupied in shSCR and sh*CDH1*-bearing mice. Data represent mean ± SD of 20 lobes from 4 mice. Data represent mean ± SD, paired t-test. **s.** Weight of xenograft glands at end-point from 16 Vector and *ZEB1*-overexpressing MCF-7 xenograft glands. Data represent mean ± SD, unpaired Student’s t-test**. t.** Relative fold change in ex vivo organ bioluminescence from 4 mice, each. Data represent mean ± SD, Multiple t-tests. **u.** Weight of xenograft glands at end-point from 28 shSCR and 34 sh*CDH1* T99 xenograft glands (8 and 9 mice, respectively), and 43 shSCR and 47 sh*CDH1* METS15 xenograft glands (9 mice for both). Data represent mean ± SD, unpaired Student’s t-test. **v.** Relative fold change in ex vivo organ bioluminescence in 8 shSCR and 9 sh*CDH1* T99 xenografts-bearing mice, and 9 shSCR and 9 sh*CDH1* METS15 xenografts-bearing mice, Data represent mean ± SD, multiple t-tests. **w.** Mammary gland weight at end-point of T99 Vector and *ZEB1* intraductal xenografts. Data represent mean ± SD of 15 and 18 xenograft glands, respectively. **x.** Relative fold change in ex vivo organ bioluminescence in 4 Vector and 5 *ZEB1*-overexpressing T99 xenografts-bearing mice, respectively. Multiple t-tests. *, **, ***, ***, and n.s represent p<0.05, 0.01, 0.001, 0.0001, and not significant, respectively.

**Extended Data Fig. 5. EMP to-epithelial transition *in vitro*. a.** Ex vivo culture of GFP+ murine cells and RFP+ MCF7 lung DTCs after lung dissociation 3 days and 2 months after plating and **b.** primary tumor cells 3 and 10 days after plating. Scale bar, 200 µm. **b. c.** Relative mRNA expression levels of selected genes in *in vitro* cultured primary, lung, and brain MCF-7 metastatic cells mice normalized to *GAPDH.* Data represent mean ± SD from n=3 control, 4 lung, and 2 brain replicates. **d.** Summary of the relative expression levels of *CDH1* (left) and *ZEB1* (right) in intraductally grafted MCF7 cells either *in situ*, invasive, DTCs, or after 2 months of *in vitro* culture. Data represent mean ± SD. EMP: Epithelial-Mesenchymal Plasticity; MET: EMP-Epithelial Transition. **e.** Western blot validation and quantification of E-cad overexpression in PLX304 (vector) and PLX*CDH1* MCF-7 cells and **f.** E-cad induction by 2 µg/ml DOX in PLIX403 (vector) and PLIX403-*CDH1* MCF-7 cells *in vitro*.

## Acknowledgements

We thank S. Egan for critical comments and H. Quinn and P. den Hollanader for careful reading of the manuscript, L. Battista for technical assistance, J. Dessimoz at the EPFL histology core facility, T. Laroche at the EPFL bioimaging and optics platform (BIOP), and B. Mangeat at the EPFL gene expression core facility (GECF) for technical assistance. We extend our gratitude to the patients who participated in our study. P.A. was supported by SNF 310030_179163/1 Exploring key steps of the metastatic cascade in ER+ breast cancer in vivo, Y.Z. by KFS-4738-02-2019-R Different facets of estrogen receptor alpha (ER) signalling during ER+ breast carcinogenesis, and G.S. by Biltema ISREC Foundation Cancera Stiftelsen, Mats Paulssons Stiftelse, and Stiftelsen Stefan Paulssons Cancerfond.

## Author contributions

Conceptualization P.A., Y. Z., C.B.; Formal Analysis P.A., Y. Z., C.S.; Investigation P.A.,Y. Z., C.S., G.S.; Resources S.M., Writing P. A., C.B; Funding Acquisition C.B.

## Materials and Methods

### Clinical samples

The Commission cantonale d’éthique de la recherche sur l’être humain (CER-VD 38/15, PB_2016-01185 (38/15)) approved this study and informed consent was obtained from all subjects. Tumor tissue was mechanically and enzymatically digested using parallel razor blades and collagenase, as previously described (Fiche et al., 2019; Sflomos et al., 2016). Effusion samples (pleura or ascites) were centrifuged at 2,500 RPM at 25°C for 10 minutes. Pellets were rinsed with phosphate buffered saline (PBS), 2% calf serum (CS), and erythrocyte-lysed using red blood cell lysis buffer (Sigma, R7757) for 5 minutes, then diluted in PBS 2% CS, and centrifuged again. Patient-derived tumor cells were transduced with either ffLuc2/Turbo-GFP or ffLuc2/Turbo-RFP (GEG-tech) overnight in low attachment culture plates (Corning® Costar® Ultra-Low Attachment) in a humidified incubator (37°C, 5% CO_2_, and 5% O_2_).

### Animal Experiments

All mice were maintained and handled according to Swiss guidelines for animal safety with a 12-h-light-12-h-dark cycle, controlled temperature and food and water *ad libitum*. Experiments were performed in accordance with protocol VD1865.5 approved by the Service de la Consommation et des Affaires Vétérinaires, Canton de Vaud, Switzerland. NOD.Cg-*Prkdc^scid^ Il2rg^tm1Wjl^*/SzJ (NSG) and NOD.Cg-*Prkdc^scid^ Il2rg^tm1Wjl^* Tg(CAG-EGFP)1Osb/SzJ (NSG-EGFP) mice were purchased from Charles River and The Jackson Laboratory. For the induction of E-cad, doxycycline (0.62g/kg of food, SAFE 150 SP-25 www.safe-diets.com) was administered in the diet.

### Intraductal injections and re-transplantation

Mice were anesthetized by intraperitoneal injection of 200 µl of 10 mg/kg xylazine and 90 mg/kg ketamine (Graeub). Intraductal injections of single-cell suspensions were performed as previously detailed (Behbod et al., 2009; Sflomos et al., 2016). Intraductal xenografts of cell lines and patient-derived cells were generated by injecting 1×10^5^ and 2×10^5^ cells, respectively, into the teats of 8-16-week-old NSG or NSG-EGF female mice, and grown for 4-6 months. For re-transplantation of patient-derived cells, mammary glands were collected on ice-cold 1X PBS, dissociated using parallel razor blades, and enzymatically digested using the tumor dissociation kit (Miltenyi Biotec) to generate single cells (human and mouse). To enrich for human cells, mouse cells were depleted using the mouse cell depletion kit (Miltenyi Biotec) according to the manufacturer’s instructions. Cells were then counted and 2×10^5^ cells were intraductally injected.

### Cell culture, cloning, and cell growth

ER+ breast cancer cell lines MCF7 and T47D as well as ER- breast cancer cell lines BT20 and HCC1806 were purchased from American Type Culture Collection (ATCC). MCF-7, T47D, BT20, and HCC1806 were maintained in Dulbecco’s modified Eagle’s medium (DMEM) medium (cat# 31966, Gibco) supplemented with 10% FCS (cat# 10270-106, Thermo Fisher Scientific Inc.) and penicillin/streptomycin (P/S, cat# 15070-063, Thermo Fisher Scientific Inc.). All cells were grown in a humidified incubator (37°C, 5% CO_2_), and were passaged when confluency reached 80%. Cells used in vivo were transduced with either ffLuc2/Turbo-GFP or ffLuc2/Turbo-RFP, and selected for the brightest fluorescent subpopulation by FACS sorting. shFF3 and sh*ZEB1*, PstByGFP and PstByZeb1 were kind gifts from Sendurai Mani (MD Anderson). Inducible E-cad vector was generated by annealing the ORF-*CDH1* (Clone ID:IOH46767, Invitrogen) cassette into an inducible backbone (pLIX403, cat# 41395, Addgene, https://www.addgene.org/41395/) using the Gateway cloning strategy (cat# 12535, Invitrogen). pLKO.1 puro shRNA E-cad was a gift from R.A. Weinberg (Addgene plasmid # 18801; http://n2t.net/addgene:18801 ; RRID:Addgene_18801). For loss of function studies, pLKO.1 puro and pLKO.1 sh*CDH1* (Onder et al., 2008) were purchased from Addgene. pLL3.7m-Clover-Geminin(1-110)-IRES-mKO2-Cdt(30-120) was a gift from Michael Lin (Addgene plasmid # 83841 ; http://n2t.net/addgene:83841 ; RRID:Addgene_83841). Lentiviruses were produced and purified as previously described (Barde et al., 2010).

### Tumor growth and metastasis analysis

Tumor growth was monitored every other week by in vivo imaging system (IVIS, Caliper Life Sciences). Briefly, 12 minutes after intraperitoneal administration of 150 mg/kg luciferin (cat# L-8220, Biosynth AG), mice were anesthetized in an induction chamber (O_2_ and 2% isoflurane) and placed inside the machine. Images were acquired and analyzed using Living Image Software (Caliper Life Sciences, Inc.). For ex vivo bioluminescence measurements, mice were first injected with 300 mg/kg luciferin for 7 minutes, then injected with 1 ml of 10 mg/kg xylazine and 90 mg/kg ketamine. Resected organs were then imaged 20 minutes after luciferin injection. Mammary glands were fixed in 4% paraformaldehyde (cat# 0335.3, Carl Roth) overnight at 4°C or flash-frozen in liquid nitrogen for RNA or protein extraction.

### Immunohistochemistry and passive clarity

Histological staining was performed as detailed previously (Ataca et al., 2020). For passive clarity, tumor-bearing mammary glands, lungs, and brain were embedded in 5% agarose (w/v, 1X PBS, cat# 16500500, Invitrogen), left at room temperature to solidify and agarose cubes then were removed from the plastic container and mounted for Vibratome sectioning (Leica VT1200 S) using glue (cat# 14460, Ted Pella, Inc.). The buffer tray was then filled with cold PBS. Blade travel speed was 0.8 mm/sec and minimum thickness of sections was 0.5 mm. A4P0 hydrogel solution was used for passive clarity (Lloyd-Lewis et al., 2016; Yang et al., 2014) to remove endogenous fluorescence for subsequent antibody labelling. To preserve endogenous tissue fluorescence, Rapiclear 1.52 solution (RC, SunJin Lab, https://www.sunjinlab.com/) was used according to the manufacturer’s instructions with some modifications. Tissues were permeabilized in 2% v/v Triton X-100 (Sigma T8787) PBS overnight on a gentle rotor shaker at room temperature. Rapiclear 1.52 was pre-warmed at 37°C and 2 ml were added on top of the tissues for 1 hr. Tissues were mounted between 2 coverslips using RC 1.52 and iSpacers for image acquisition and long-term storage. Z-stacks were acquired using a Zeiss LSM700 confocal microscope and 3D-reconstructed using Imaris bitplane image analysis software. The list of primary and secondary antibodies is provided in Supplementary Table 4.

### Immunohistochemistry analysis

For analysis of primary tumor staining, outlines of human cells were drawn manually with “Freehand selections” and a built-in function “waitForUser” for each image to exclude mouse cells, followed by ROIs creation. Channels were then binarized using “Huang” thresholding algorithm and images were processed with watershed segmentation to split closely touching cells and then denoised by a Minimum Filter with radius of 1 pixel. Cell numbers were quantified using the “analyse particles” function of Fiji (Size 0.01-infinity pixel^2^, circularity 0-1). Finally, channels were saved and the cell quantifications were output as a table for further data analysis. For analysis of metastatic breast cancer cells in lungs, human cell nucleus and Ki67+ cells were counted manually.

For analysis of the FUCCI reporter in primary tumors, images were analysed every 8 slices from the original Z-stack images to meet the criteria of the Nyquist–Shannon sampling theorem. Green and red channels of interest were binarized using the “Li” thresholding algorithm that was chosen based on the visual inspection of output images. Images were processed with watershed segmentation to split closely touching cells and then denoised by a Minimum Filter with radius of 1 pixel. The masks of the two channels were used to generate a third mask using the “AND” operation from “Image Calculator” to indicate double positive cells. Cell numbers were quantified using the “analyse particles” function of Fiji for all 3 masks (size 1-infinity pixel^2^, circularity 0-1) and the outlines of quantified cells were recorded in another three masks (Show = outlines). Finally, the six masks (two channels of interest, one from the AND operation and their 3 masks showing outlines) were merged and saved for visualization and quality inspection. For analysis of FUCCI reporter in lungs, green, red and double positive cells were quantified manually. The percentages of cycling and non-cycling cells were quantified using the following equations:

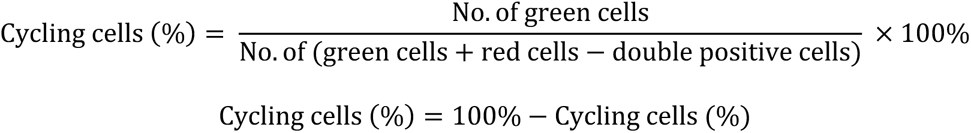

For the analysis of mesenchymal-like cells in primary tumors, the blue channel of interest (DAPI) was binarized using the “Percentile” thresholding algorithm for MCF7 and T47D and “Moments” for METS15 tumors based on the visual inspection of output images. Images were processed with watershed segmentation to split closely touching cells, followed by cell aspect ratios (ARs) quantification using the “analyse particles” function of Fiji (Size 50-infinity pixel^2^, circularity 0-1). Finally, the mask for the blue channel was saved and the cell quantifications were output as a table for further data analysis. For analysis of mesenchymal-like cells in lungs, major and minor axis were drawn manually on each breast tumor cell and then measured to quantify ARs. At least 300 metastatic cells in lungs from each mouse of 3 mice in total were measured. Mesenchymal-like cells were defined as those with AR > 1.6 based on the following equation:

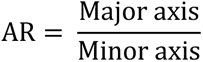

For the analysis of fluorescence intensity, corrected total cell fluorescence (CTCF) was calculated by manually drawing an outline around the border of the cells using the drawing/selection tools in ImageJ, and a close-by region with no cells to serve as a measure of background noise. Measurements were generated by ImageJ and CTCF was calculated using the following formula:

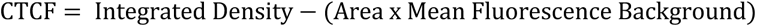

### Whole mounts

Fluorescence images of whole mammary glands, lungs, and brains were acquired with a LEICA M205FA fluorescence stereo microscope equipped with a Leica DFC 340FX camera.

### RNA extraction and RT-PCR

Cell pellets, tumor-bearing mammary glands, and microdissected lung, liver, and brain micrometastases were homogenized with TRIzol reagent (Invitrogen), total RNA was isolated with miRNeasy Mini Kit (Qiagen), cDNA was synthesized with random p(dN)_6_ primers (Roche) and MMLV reverse transcriptase (Invitrogen). RT-PCR analysis in triplicates was performed with SYBR Green FastMix (Quanta) reaction mix. Supplementary Table 5 provides list of primers.

### Immunoblotting

Cell pellets and tumor-bearing mammary glands were homogenized in 1% SDS solution containing protease (cat# 11836153001, Roche AG) and phosphatase inhibitor cocktail (cat# 04906845001, PhosStop, Roche, AG), followed by 20 min centrifugation at 4°C (14,000 x g). Protein-containing supernatant was then quantified and normalized to a reference protein using the Image Studio Lite version 5.2 (LI-COR®). SDS-polyacrylamide and transfer were then performed as previously detailed (Sflomos et al., 2016). Membranes were revealed with ECL or WesternSure PREMIUM Chemiluminescent Substrate (cat# 926-95010, LI-COR®). The primary and secondary antibody list can be found in Supplementary Table 4.

### Statistics

Statistical analysis was performed using GraphPad Prism 8 (San Diego, California, USA, www.graphpad.com). Statistical tests are indicated in the figure legends. For growth curves, two-way ANOVA with multiple comparisons was used, for contralateral intraductal grafts, paired Student’s t-test was performed, and for individual grafts, unpaired Student’s t-test were performed, while testing for normality (Shapiro-Wilk test). Non-parametric statistical tests were used when Kolmogorov-Smirnov and Shapiro-Wilk normality tests failed to show a normal distribution.

