## Supplementary figures and images for "Epithelial-mesenchymal plasticity determines estrogen receptor positive (ER+) breast cancer dormancy and reacquisition of an epithelial state drives awakening"

### Supplemental Figures

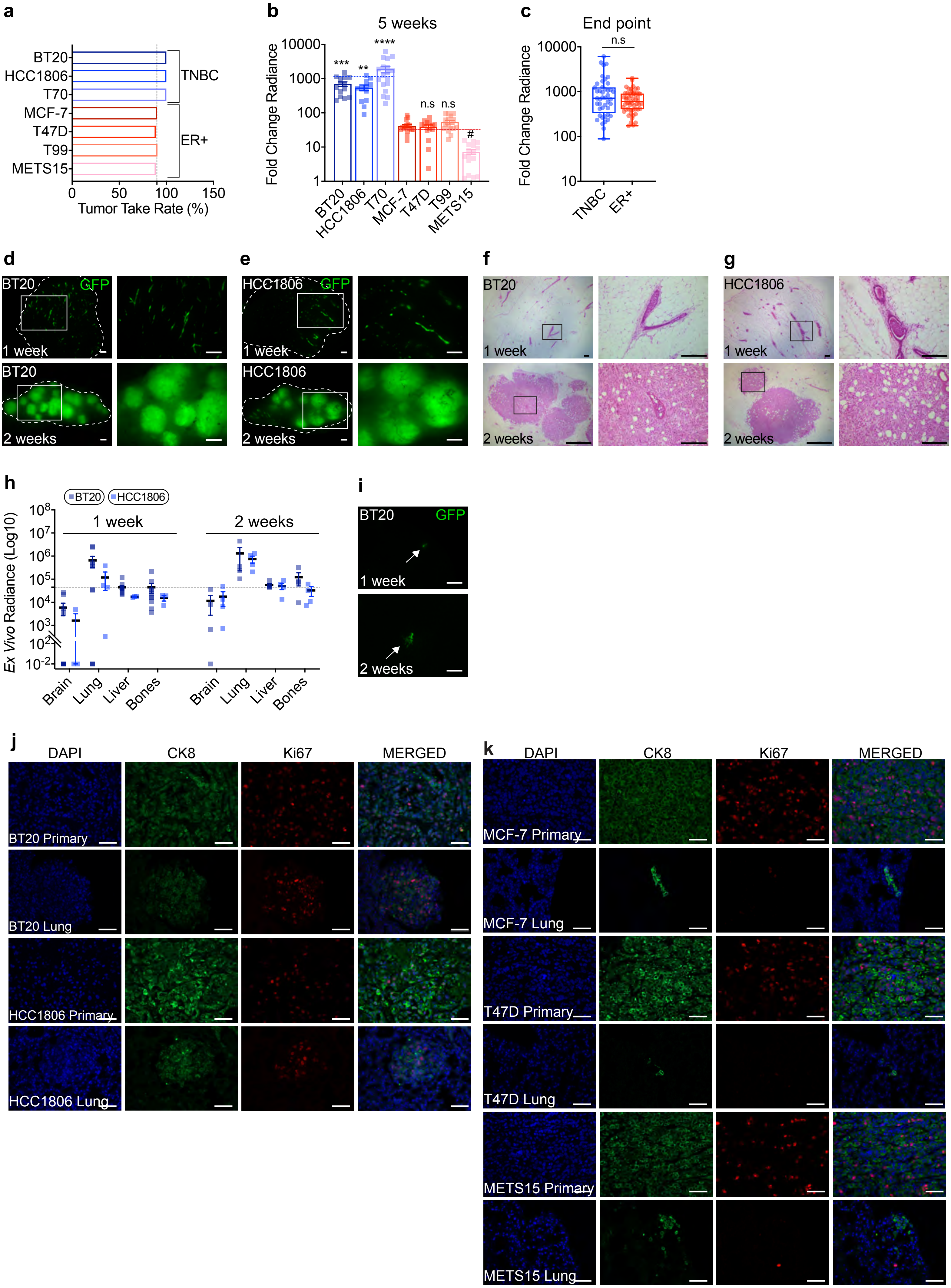

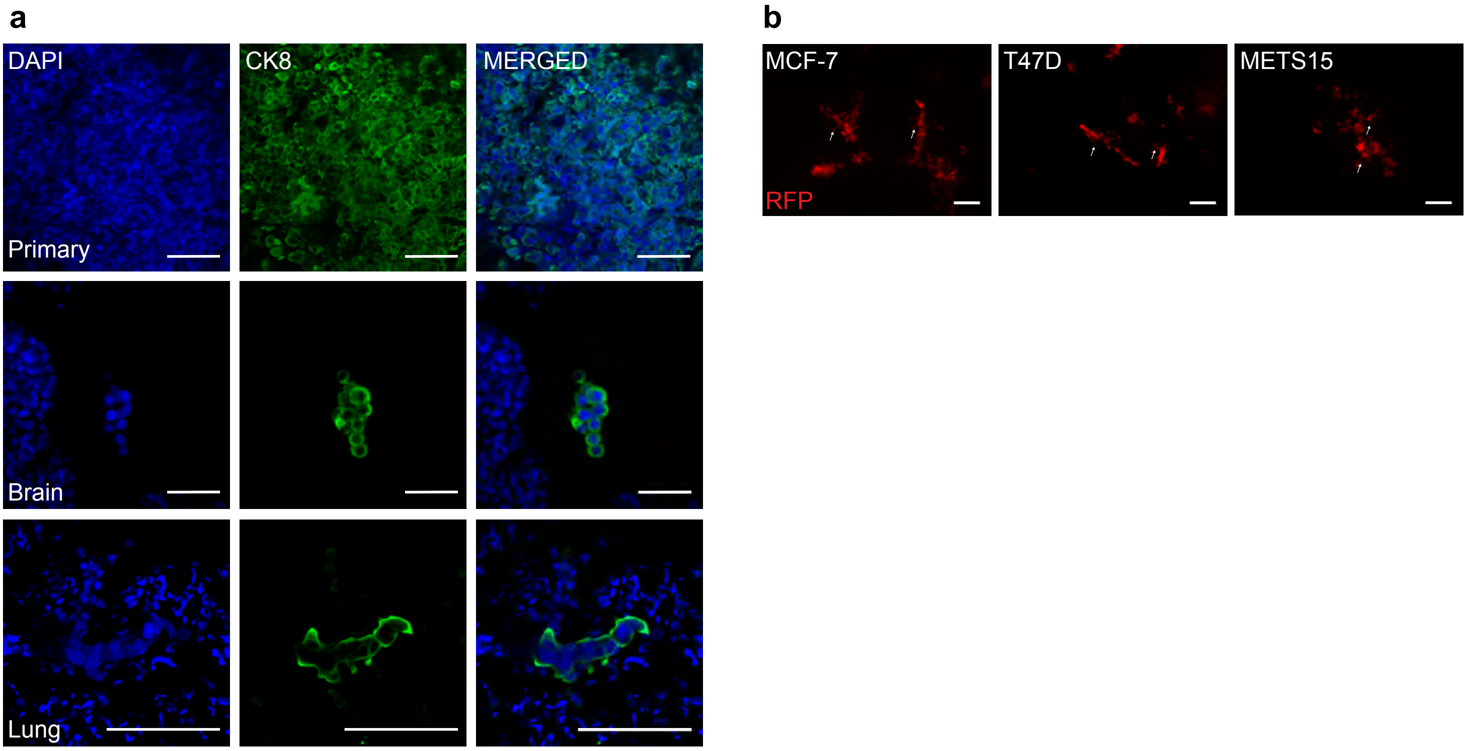

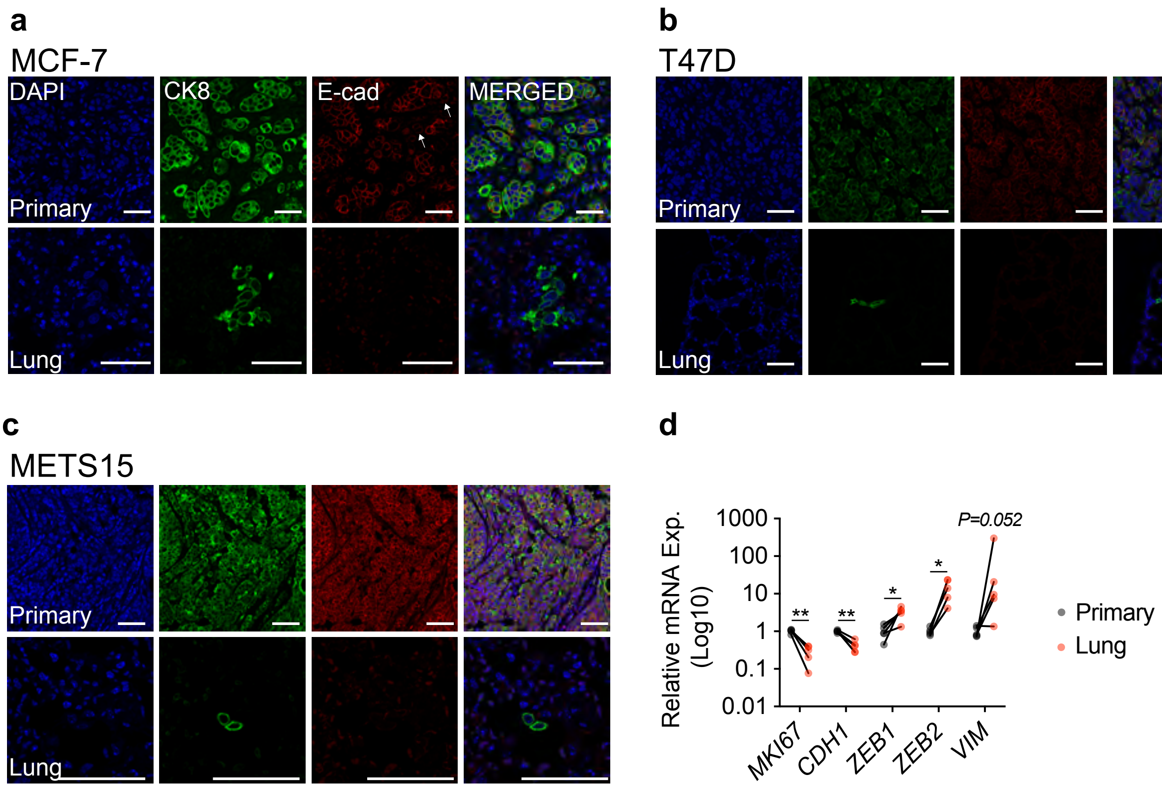

Aouad et al.

Extended Data Figure 4

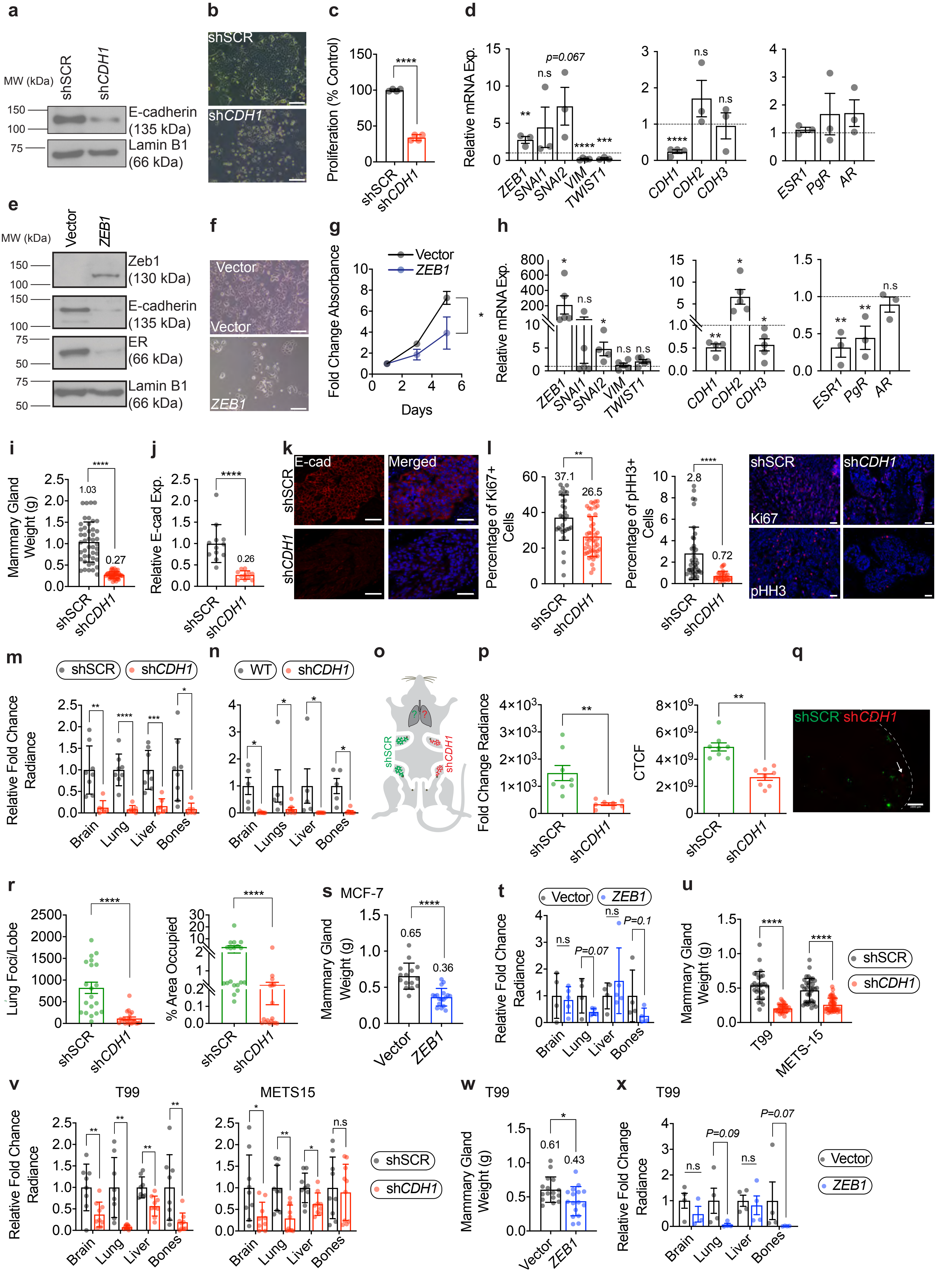

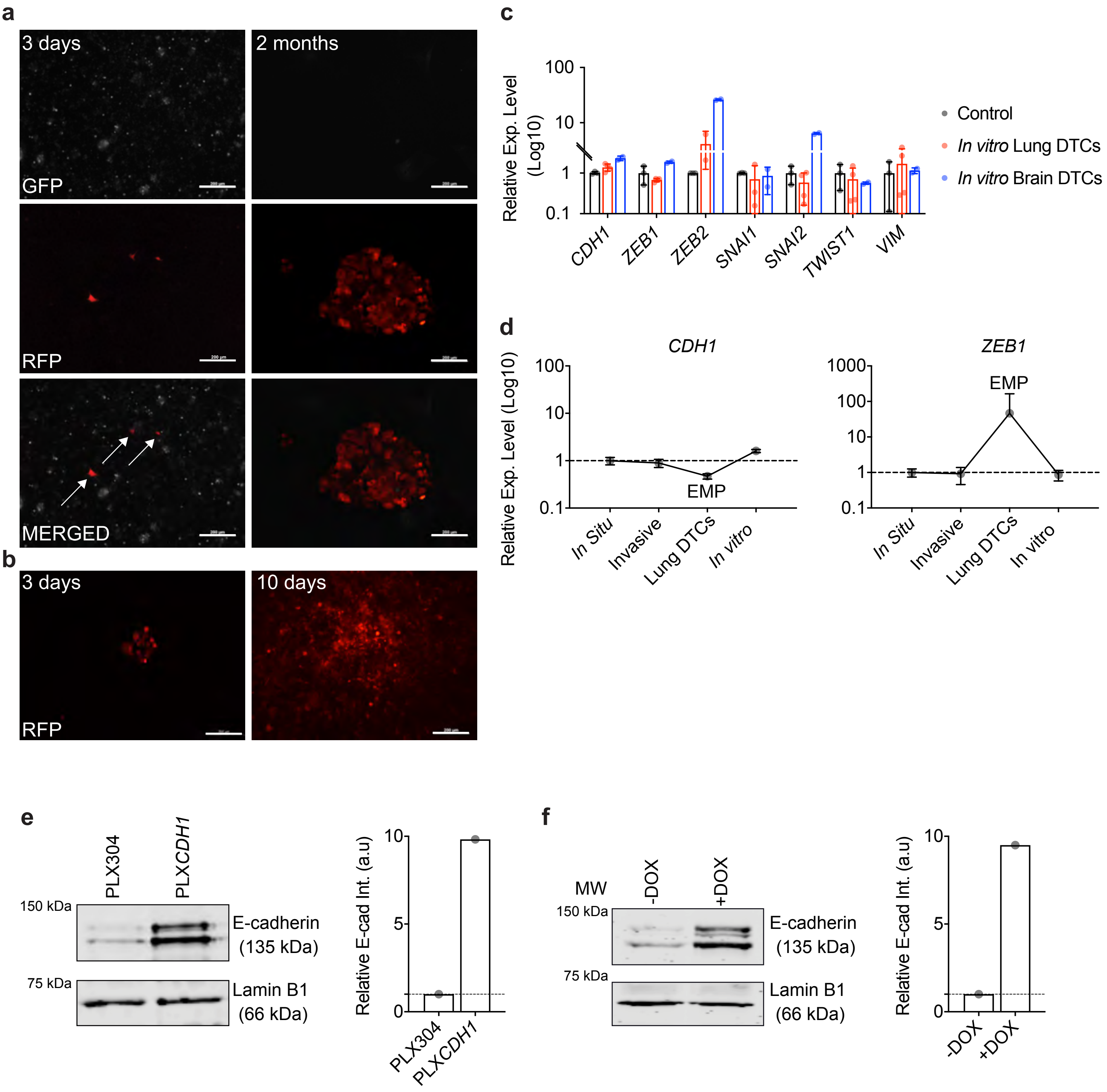
