## Supplemental Tables for "Epithelial-mesenchymal plasticity determines estrogen receptor positive (ER+) breast cancer dormancy and reacquisition of an epithelial state drives awakening"

| Cell line | Age | Source | ER | PR | HER2 | Subtype | in vitro Passage* |
|---|---|---|---|---|---|---|---|
| MCF-7 | 69 | Pleural Effusion | + | + | - | Luminal | 7-30 |
| T47D | 54 | Pleural Effusion | + | +++ | - | Luminal | 6-8 |
| BT20 | 74 | Primary Tumor | - | - | - | Basal-like | 6-11 |
| HCC1806 | 60 | Primary Tumor | - | - | - | Basal-like | 20-23 |

Supplemental Table 1. Table describing the cell lines used in this study.
+: Positive
-: Negative
*in vitro Passage after obtention from ATCC

| PDX | Age | Tumor Type | ER% | PR% | HER2% | Ki67% |
|---|---|---|---|---|---|---|
| T99 | 57 | NST, Primary | 57 | 70 | - | 20 |
| METS15 | 59 | NST, Ascites | 90 | 100 | - | N/A |
| T70 | 39 | NST, Primary | - | - | - | >90 |

| PDX | Treatment | PDX Generation |
|---|---|---|
| T99 | Untreated | 6 to 8 |
| METS15 | Chemo, AI, FULV | 1 to 4 |
| T70 | Untreated | 6 to 8 |

Supplemental Table 2. Table describing patient-derived primary cells characteristics and the generated PDXs. NST: no special type; -: negative; N/A: not applicable; Chemo: chemotherapy; AI: aromatase inhibitors; FULV: fulvestrant.

| | Primary | Lung |
|---|---|---|
| MCF-7 | 38,847 | 1,909 |
| T47D | 43,162 | 548 |
| METS15 | 62647 | 1087 |
| BT20 | 30,459 | 2,593 |
| HCC1806 | 11,619 | 2,099 |

Supplemental Table 3. Total number of Ki67-positive cells in matched primary and lung metastases in ER+ and TNBC intraductal xenografts.

| Antibody | Source | Identifier | Clone | Usage | Dilution |
|---|---|---|---|---|---|
| CK8 | BioLegend | MMS-162P-250 | 1E8 | IF | 1 in 400 |
| DsRed | MBL Int. | PM005 | N/A | IF | 1 in 500 |
| E-cadherin | Cell Signaling | 3195S | 24E10 | WB, IF | 1 in 100 |
| ER | Ventana Medical Systems | 790-4324 | SP-1 | WB | 1 in 200 |
| GFP | Santa Cruz | sc-9996 | B-2 | IF | 1 in 500 |
| Ki67 | ThermoFisher Scientific | MA5-14520 | SP-6 | IF | 1 in100 |
| Lamin B1 | Abcam | AB16048 | N/A | WB | 1 in 1000 |
| pHH3 (ser10) | Millipore | 06-570 | 3H10 | IF | 1 in 500 |
| p27 Kip1 | Cell Signaling | 3686 | D69C12 | IF | 1 in 100 |
| Zeb1 | Novus Biologicals | NBP1-05987 | N/A | WB | 1 in 500 |
| Mouse Alexa 488 | ThermoFisher Scientific | A-11029 | N/A | IF | 1 in 500 |
| Mouse Alexa 568 | ThermoFisher Scientific | A-10037 | N/A | IF | 1 in 500 |
| Mouse Alexa 647 | ThermoFisher Scientific | A-31571 | N/A | IF | 1 in 500 |
| Rabbit Alexa 488 | ThermoFisher Scientific | A-21206 | N/A | IF | 1 in 500 |
| Rabbit Alexa 568 | ThermoFisher Scientific | A-10042 | N/A | IF | 1 in 500 |
| Rabbit Alexa 647 | ThermoFisher Scientific | A-31573 | N/A | IF | 1 in 500 |
| Rat Alexa 647 | ThermoFisher Scientific | A-21247 | N/A | IF | 1 in 500 |

Supplemental Table 4. List of antibodies used in this study. IF: immunofluorescence, WB: Western blot.

| Gene | Forward Primer | Reverse Primer | Reference |
|---|---|---|---|
| *CDH1* | 5'-TGC CCA GAA AAT GAA AAA GG-3' | 5'-GTG ATA GTG GCA ATG CGT TC-3' | 1 |
| *CDH2* | 5'-ACA GTG GCC ACC TAC AAA GG-3' | 5'-CCG AGA TGG GGT TGA TAA TG-3' | 1 |
| *CDH3* | 5'-ATC ATC GTG ACC GAC CAG AAT-3' | 5'-GAC TCC CTC TAA GAC ACT CCC-3' | N/A |
| *ZEB1* | 5'-ACC TGC CAA CAG ACC AGA CAG TGT-3' | 5'-CCT GAC CTT CAG GCC CCA GGA-3' | N/A |
| *ZEB2* | 5'-AGC CAC GAT CCA GAC CGC AA-3' | 5'-GCT GTG TCA CTG CGC TGA AGG T-3' | N/A |
| | 5'-AAT GCA CAG AGT GTG GCA AGG C-3' | 5'-CTG CTG ATG TGC GAA CTG TAG G-3' | N/A |
| *SNAI1* | 5'-GGA GTC CGC AGT CTT ACG AG-3' | 5'-TCT GAA GAA CCT GGT AGA GG-3' | 1 |
| *SNAI2* | 5'-CTG GGC GCC CTG AAC ATG CAT-3' | 5'-GGC TTC TCC CCC GTG TGA GTT CTA-3' | 2 |
| *VIM* | 5'- GCA AAG ATT CCA CTT TGC GT-3' | 5'-GAA ATT GCA GGA GGA GAT GC-3' | 3 |
| | 5'-GAG AAC TTT GCC GTT GAA GC-3' | 5'-GCT TCC TGT AGG TGG CAA TC-3' | N/A |
| *TWIST1* | 5'-CAT CCT CAC ACC TCT GCA TT-3' | 5'-GGC CAG TTT GAT CCC AGT AT-3' | 1 |
| *GAPDH* | 5'-AGG GCT GCT TTT AAC TCT GCT-3' | 5'-CCC CAC TTG ATT TTG GAG GGA-3' | N/A |
| *HPRT* | 5'-TAG AAG GCC TTG TGC TCA CC-3' | 5'-TCT GCT CTG ACT TTA GCA CCT G-3' | N/A |
| *MKI67* | 5'-CGG ACT TTG GGT GCG ACT T-3' | 5'-GTC GAC CCC GCT CCT TTT-3' | N/A |
| *TBP* | 5'-TAG AAG GCC TTG TGC TCA CC-3' | 5'-TCT GCT CTG ACT TTA GCA CCT G-3' | N/A |
| *ESR1* | 5'-GGA GAT CTT CGA CAT GCT GC-3' | 5'-GCC ATC AGG TGG ATC AAA GT-3' | N/A |
| *PgR* | 5'-AAA CTG CCC AGC ATG TCG TCT-3' | 5'-GCT CTG GTT AGG AAG GCC CA-3' | N/A |
| *AR* | 5'-GAG AAC TTT GCC GTT GAA GC-3' | 5'-GCT TCC TGT AGG TGG CAA TC-3' | N/A |

Supplemental Table 5. List of primers used in this study.
